# Context modulates brain state dynamics and behavioral responses during narrative comprehension

**DOI:** 10.1101/2025.04.05.647323

**Authors:** Yibei Chen, Zaid Zada, Samuel A. Nastase, F. Gregory Ashby, Satrajit S. Ghosh

## Abstract

Narrative comprehension is inherently context-sensitive, yet the brain and cognitive mechanisms by which brief contextual priming shapes story interpretation remain unclear. Using hidden Markov modeling (HMM) of fMRI data, we identified dynamic brain states as participants listened to an ambiguous spoken story under two distinct narrative contexts (affair vs. paranoia). We identified recurrent states involving auditory, language, and default mode network (DMN) regions that were expressed across both groups, as well as additional states characterized by recruitment of multiple-demand network (MDN) systems, including control, dorsal attention, and salience networks. Bayesian mixed-effects modeling revealed that contextual framing modulated how specific linguistic and character-related features influenced the probability of occupying these states. Complementary behavioral data showed parallel context-sensitive modulation of participants’ moment-to-moment interpretive judgments. Together, these findings suggest that contextual priming influences narrative comprehension through subtle, feature-dependent adjustments in the engagement of DMN- and MDN-related brain states during naturalistic story listening.

## Introduction

Narrative comprehension involves complex interactions among prior knowledge, immediate contextual expectations, and the content (Botch & Finn, 2024; Mar & Oatley, 2008; Nastase et al., 2021; Willems et al., 2020). Recent neuroimaging research demonstrates significant variability in how individuals process identical narrative stimuli, primarily driven by stable personal traits such as empathy, political beliefs, and cognitive abilities. This variability results in distinct patterns of brain activity and synchronization (Coderre & Cohn, 2023; de Bruin et al., 2023; Johns et al., 2018; Nijhof & Willems, 2015). In addition to these stable individual differences, transient manipulations of expectations or interpretation profoundly impact how narrative content is processed, highlighting the brain’s sensitivity to context (Yeshurun et al., 2017).

Research in psychology and linguistics highlights the crucial role of context in understanding narratives. Contextual cues, such as background knowledge or primed information, help people construct coherent mental representations, improve comprehension, and enhance recall of narrative details (Bransford & Johnson, 1972; van Kesteren et al., 2013; Zwaan & Radvansky, 1998). These longer narratives allow context to accumulate gradually, shaping interpretation as the story progresses and enabling the construction of shared mental models across listeners.

Neuroimaging studies have shown that naturalistic narratives (e.g., audiostories or movies), which unfold over time, recruit default mode network regions that are typically less engaged by highly controlled lab stimuli that are shorter and decontextualized (Baldassano et al., 2017; Ben-Yakov et al., 2012; Geerligs et al., 2022; Lerner et al., 2011; Yeshurun et al., 2021). Recent theoretical work proposes that the DMN supports the construction and maintenance of mental models more generally, summarizing high-dimensional experiences into lower-dimensional, situation-level representations (Barnett & Bellana, 2025). Converging network-level evidence suggests that externally presented narratives are processed hierarchically along the cortical gradient, from primary sensory cortices (e.g., auditory) to the language network and ultimately to the DMN (Gordon et al., 2020). Through this hierarchical progression, auditory, language, and DMN regions collectively support the integration of incoming speech into a coherent situation model, forming a dominant processing mode that sustains narrative comprehension over extended timescales.

When individuals receive similar contextual information, whether through common prior knowledge or priming, their cognitive and emotional responses tend to align, producing synchronized activity across brains (de Bruin et al., 2023; Lahnakoski et al., 2014; Nguyen et al., 2019). Conversely, when people are primed differently or bring distinct prior experiences to the same narrative, they may interpret it in diverging ways, resulting in idiosyncratic patterns of brain activity (Jacoby & Fedorenko, 2020; Yeshurun et al., 2017). These interpretive processes rely on integrating incoming information with existing mental models and are thought to emerge from coordinated activity across multiple large-scale brain networks, rather than being localized to any single region (Barrett, 2022; McIntosh, 2004; Song et al., 2023).

These processes often require updating mental models in response to shifts in narrative content, which increases cognitive demands on the system (Yang et al., 2023). Such effortful updating and ambiguity resolution reliably recruit the multiple-demand network (MDN), which involves a set of frontoparietal, dorsal attention, and salience networks engaged by a wide range of cognitively demanding tasks (Cole et al., 2013; Duncan, 2010; Hermans et al., 2014; Uddin, 2015). During narrative comprehension, these MDN components support functions such as maintaining or revising interpretive hypotheses, reorienting attention to salient cues, detecting conflict between contextual expectations and incoming information, and guiding top-down control over narrative interpretation.

Narrative comprehension is therefore not a unitary process but a dynamic interplay between multiple neural systems. Unlike traditional laboratory tasks that isolate a single function, naturalistic stories require listeners to continually shift between perceiving speech, integrating semantic information, retrieving relevant knowledge, monitoring contextual cues, and making inferences about characters and events. These shifting demands naturally recruit different large-scale networks at different times, including transitions between DMN-dominated integrative processing and MDN-dominated evaluative or attention-driven processing. Contextual priming can bias which of these processing modes is engaged at particular moments, yet the mechanisms through which such external framing shapes these evolving neural patterns remain poorly understood.

In this study, we examine how the integration of narrative input with initial contextual priming is reflected in dynamic patterns of brain activity, using the concept of “brain states.” Brain states refer to recurring patterns of coordinated activity across distributed brain regions (Liu et al., 2025; Song et al., 2021), analogous to distinct musical motifs formed by different instruments in an orchestra. Because naturalistic narratives engage different functional systems at different moments, a brain-state framework allows us to capture both the states themselves and the transitions between them, providing a window into how the brain alternates between competing or complementary processing modes over time (Shine et al., 2019; Vidaurre et al., 2017). By identifying and characterizing these recurrent states, we can assess how contextual framing influences not only which large-scale networks are engaged, but also how the brain traverses between integrative DMN-related processing and more effortful MDN-related evaluative modes as the story unfolds.

Another critical gap remains in understanding how contextual priming interacts with specific narrative features, such as character identity and other linguistic structures. During story listening, character identity is often conveyed and reinforced through character speech, especially when direct quotations are attributed to particular speakers. These attributions fundamentally shape comprehension by guiding attention, emotional engagement, and social inference (Gerrig, 1993; Jacoby & Fedorenko, 2020; Mar & Oatley, 2008). Psycholinguistic evidence consistently underscores the central role of character speech in maintaining narrative coherence, supporting mental simulation, and enabling theory-of-mind reasoning (Nieuwland & Van Berkum, 2006; Zwaan & Radvansky, 1998). Thus, character speech serves as a theoretically meaningful and empirically tractable feature for investigating how contextual priming influences narrative processing.

To investigate how narrative context shapes brain state dynamics during story comprehension, we used a naturalistic fMRI paradigm in which two participant groups listened to the same story but were primed differently beforehand (Yeshurun et al., 2017). We applied hidden Markov models (HMMs) to identify recurrent brain states, defined as temporally evolving patterns of network-level activity, across the full duration of story listening (Quinn et al., 2018; Shine et al., 2019; Taghia et al., 2018; Vidaurre et al., 2017). Based on work demonstrating that naturalistic narratives engage both DMN-supported semantic integration and MDN-supported evaluative or attention-driven processing, we expect that narrative comprehension would elicit at least two broad classes of brain states: (i) integration states reflecting coordinated activity among auditory, language, and default mode networks, and (ii) evaluative states involving control, dorsal attention, and salience networks that support ambiguity resolution and situation updating. Building on prior work showing brain sensitivity to character-level features (Alderson-Day et al., 2020; Jacoby & Fedorenko, 2020; Yarkoni et al., 2008), we further predicted that brain state dynamics would differ based on speaker identity.

We added a complementary behavioral experiment to better understand how primed context influences moment-to-moment interpretation. Our goal was to capture when listeners subjectively recognized elements of the story as aligning with their assigned context. Two separate groups of participants received the same context instructions and listened to the same story as those in the fMRI study. They were asked to press a key whenever they perceived information consistent with their contextual framing. These responses provide a time-resolved behavioral index of interpretive alignment, offering an external marker of how context interacts with narrative features over time.

## Methods

## fMRI dataset

We utilized the “prettymouth” dataset (Figure 1) (Yeshurun et al., 2017), which includes 40 participants drawn from the Narratives data collection (Nastase et al., 2021). Participants were divided into two groups (initially N = 20 per group), with both groups exposed to an adapted version of J. D. Salinger’s short story, "Pretty Mouth and Green My Eyes." The adapted version was shorter than the original and included several sentences not present in the original text. A professional actor provided the narration, resulting in a recording of 11 minutes and 32 seconds. Functional MRI data were acquired with a repetition time (TR) of 1.5 seconds. The story was preceded by 18 seconds of neutral music and 3 seconds of silence, followed by an additional 15 seconds. These segments of music and silence were excluded from all analyses.

**Figure 1.**
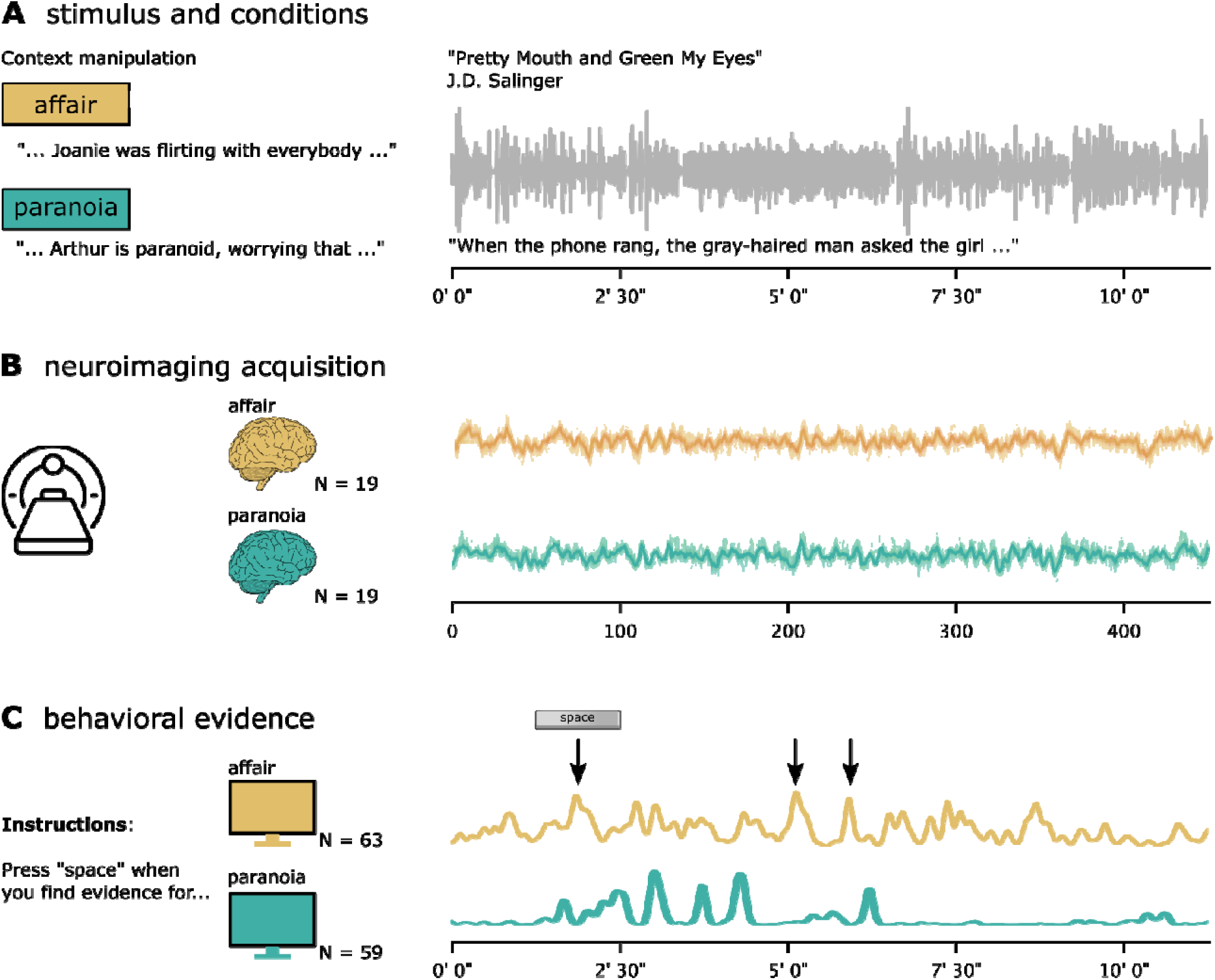
Experimental design, neuroimaging acquisition, and behavioral evidence. (**A**) Participants were randomly assigned to one of two context conditions—affair (gold) or paranoia (teal)—and read a brief prompt before listening to an 11-minute spoken story (Pretty Mouth and Green My Eyes by J.D. Salinger). This context manipulation was consistent across both neuroimaging and behavioral experiments. (**B**) The fMRI study included data from 19 participants in each context group in the final analysis. The schematic plot illustrates average fMRI time courses for each group (affair in gold, paranoia in teal), with the x-axis representing time in TRs. (**C**) In the behavioral study, we recruited two new groups of participants and asked them to press the spacebar whenever they perceived evidence supporting one interpretation or the other (affair N = 63, paranoia N = 59). The line plots depict the average frequency of button presses over time within each group (i.e., agreement across all participants), with peaks corresponding to moments of perceived narrative support for each interpretation.

The narrative describes a phone conversation between two friends, Arthur and Lee. Arthur, who has just returned home from a party after losing track of his wife Joanie, calls Lee to express his concerns about her whereabouts. Lee is at home with a woman beside him, whose identity remains ambiguous: she may or may not be Joanie. Before listening to the story, each participant group received one of two different contextual prompts: one group was informed that Arthur was paranoid and his suspicions were unfounded (“paranoia” context), while the other group was told that the woman was indeed Joanie, Arthur’s wife, and that Lee and Joanie had been involved in an ongoing affair for over a year (“affair” context). Yeshurun et al. (2017) and Nastase et al. (2021) describe the experimental paradigm and fMRI data acquisition parameters.

### fMRI data processing

fMRI data preprocessing was conducted using fMRIPrep (version 24.0.1) (Esteban et al., 2019) via the BIDS App Bootstrap (Zhao et al., 2024), with detailed processing steps provided in the supplementary material. Custom post-processing steps optimized for narrative listening analyses were applied after initial preprocessing. Specifically, we implemented spatial smoothing (6 mm full-width half-maximum) to balance noise reduction and preservation of spatial activation patterns, performed detrending to mitigate scanner drift, standardized (z-scored) the time-series signals across time points within each voxel within each subject, and regressed out nuisance signals related to head motion and physiological noise using motion parameters and anatomical CompCor regressors. All post-processing steps were carried out using Nilearn; more details are provided in the supplementary material and our GitHub repository (see the Code Availability section). Post-processed data were further extracted using the Schaefer et al. (2018) parcellation (1000 parcels) with 17 networks (Kong et al., 2021). Following recommendations by Nastase et al. (2021), two participants were excluded from further analysis due to data quality concerns, resulting in a final sample size of N = 19 per group.

### Behavioral data

We collected an additional behavioral dataset under two different tasks to assess the evidence in the stimulus supporting each narrative context over time. Behavioral data were collected from 128 participants recruited via Prolific (www.prolific.com). Participants were classified into two experimental groups: an affair group (n = 63) and a paranoia group (n = 59). Demographic details indicated the sample comprised 62 males, 57 females, and three individuals identifying as non-binary or third gender. The age distribution included participants aged 18–24 years (n = 18), 25–34 years (n = 46), 35–44 years (n = 29), 45–54 years (n = 12), 55–64 years (n = 14), and 65 years or older (n = 3). These participants are a separate sample from those included in the fMRI experiment.

Data were collected through an online experiment developed using PsychoJS scripts derived from the PsychoPy builder (PsychoPy3, version 2023.2.0), hosted on Pavlovia (https://pavlovia.org/). Participants initially provided informed consent via Qualtrics (https://www.qualtrics.com/) before being randomly assigned to one of two context conditions (affair versus paranoia). Participants in each group received the same prompts presented to the fMRI participants prior to listening to the auditory story. Participants were asked to identify moments in the narrative where they perceived evidence for their assigned interpretation (Lee and Joanie are having an affair, or Arthur is being paranoid) by pressing the spacebar on their keyboards. Immediate visual feedback was provided, indicated by a brief green dot appearing at the center of the screen, confirming each response. After completing the task, participants were redirected to Qualtrics to complete a post-experiment questionnaire. Data from four participants were excluded from subsequent analyses due to incomplete records, resulting in a final dataset of 122 participants (Figure 1C). More detailed instructions can be found in the supplementary material.

This study was approved by the Princeton University Institutional Review Board (IRB 12201). In accordance with institutional ethical guidelines, all participants provided informed consent electronically before participation. Participants received monetary compensation consistent with university policy. All data were anonymized to ensure participant confidentiality.

### Hidden Markov model (HMM) analysis

To characterize the temporal dynamics of brain states during story listening, we employed hidden Markov models (HMMs, Figure 2A), which identify recurring patterns of brain network activity and their transitions over time (Baldassano et al., 2018; Meer et al., 2020; Vidaurre et al., 2017; Yang et al., 2023). HMMs explicitly model temporal dependencies and sequential state transitions, aligning closely with our objective of understanding how prior contextual information modulates the temporal evolution of brain states.

**Figure 2.**
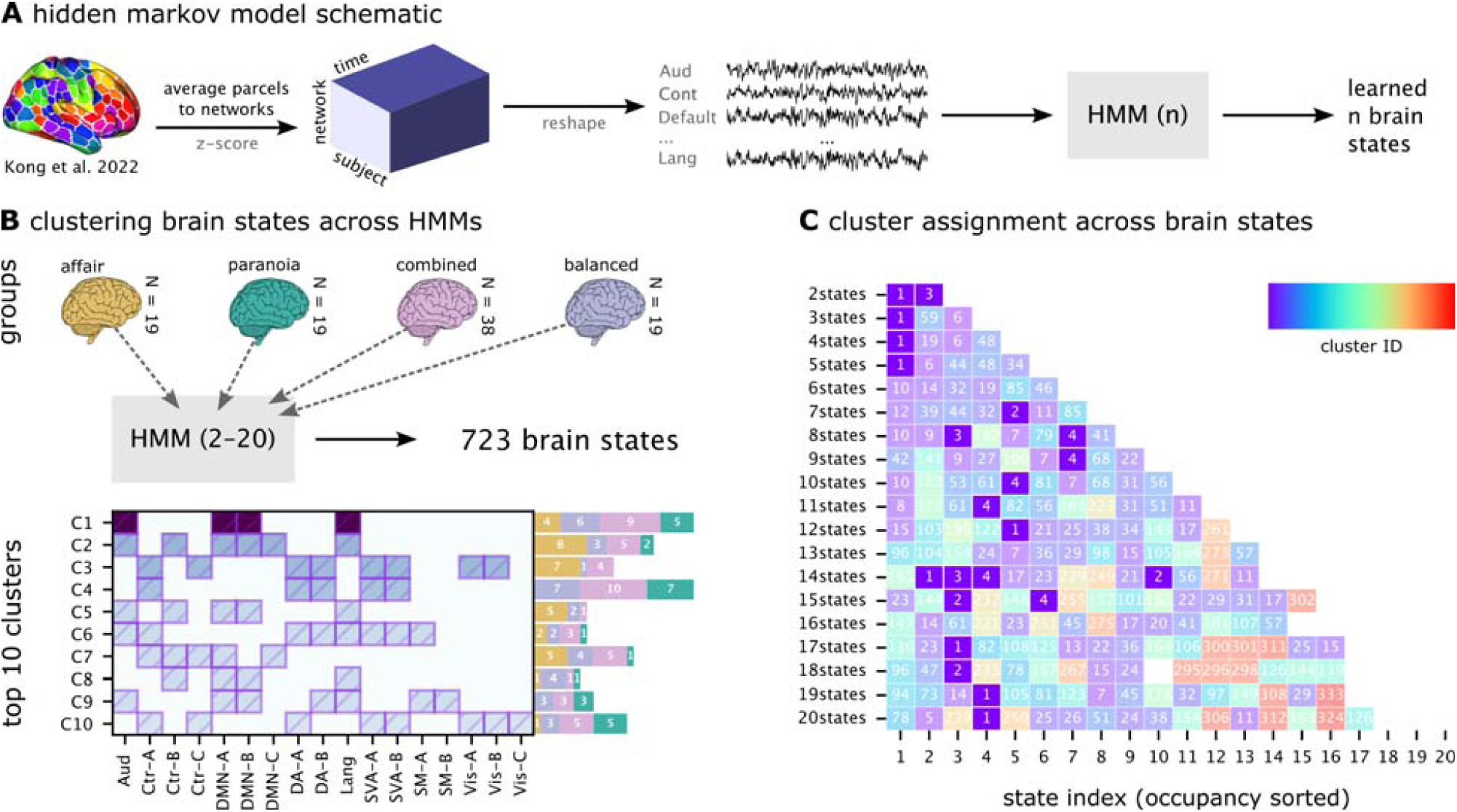
Hidden Markov models (HMMs) and clustering of brain states. (**A**) Analysis pipeline. Parcel-level fMRI time series were averaged within networks, z-scored across time, and concatenated across participants. HMMs with 2–20 states were estimated separately for four groups: *affair*, *paranoia*, *combined*, and *balanced*. (**B**) Cluster profiles and provenance. Brain states that passed quality filters (activation > 0.1, bootstrap CI width < 0.3, split-half reliability > 0.5) were pooled across models and clustered by spatial similarity (Jaccard distance, hierarchical clustering). The left panel shows the average activation profile of each resulting cluster across canonical functional networks, with shading intensity indicating the mean activation level. The bar plots to the right show the number of states from each group (affair, paranoia, combined, balanced) contributing to each cluster, illustrating cluster provenance. (**C**) Cluster assignments across model granularities. This panel visualizes how the unified cross-group clustering solution maps onto the *combined-group* HMMs. Each row corresponds to one HMM solution (2–20 states), and each column to a state within that model, ordered by state occupancy. Colors and overlaid numbers indicate the cluster assignment (1–4) of each state. Missing cells indicate states that did not pass the filtering criteria and were therefore excluded from clustering. This panel demonstrates how states from different HMM solutions converge onto a consistent set of four clusters.

To account for hemodynamic delay, the BOLD signal was shifted backward by three TRs (∼4.5 s) relative to the timing of the stimulus features (Yeshurun et al., 2017). Non-story segments (background music/silence, 24 TRs at scan onset and offset) were excluded, yielding 451 TRs for analysis. Time series were extracted from 17 functionally defined networks by first averaging voxel-wise signals within each parcel, then averaging across all parcels assigned to the same network. Each participant’s data was z-scored to normalize signal amplitudes.

We implemented Gaussian observation HMMs using the *hmmlearn* Python package, modeling brain states as multivariate Gaussian distributions with state-specific means and covariances. Transition probabilities were initialized to favor an expected dwell time of approximately 7 seconds (∼4–5 TRs). This provided a weak prior consistent with evidence that large-scale brain states during naturalistic cognition typically persist on the order of several seconds, with higher-order regions such as the default mode network exhibiting longer durations and sensory regions (e.g., auditory cortex) switching more rapidly (Vidaurre et al., 2017; Baldassano et al., 2017). The prior served to prevent implausibly fast state-switching at initialization, but transition probabilities were re-estimated during expectation–maximization, so final dwell times were determined by the data rather than fixed to 7 seconds.

To improve robustness against local optima during model fitting, we performed five independent initialization attempts per model (i.e., with the same number of states but different random seeds). For each attempt, state means were initialized by random draws from a standard normal distribution (***𝒩*(0,1)**), ensuring broad coverage of the feature space. Covariance matrices were initialized as identity matrices with a small diagonal ridge term (***є* = 10^-8^**) to maintain positive definiteness and numerical stability during optimization. Start probabilities were initialized uniformly across states, and transition priors followed the weak dwell-time prior described above. Across the five attempts, random seeds influenced both the initialization of means and the stochastic path of the EM algorithm, while covariances, start probabilities, and transition priors were held constant. The solution with the highest log-likelihood was selected for downstream analysis.

Model generalizability and stability were assessed through leave-one-subject-out cross-validation (LOOCV). For each fold, a group-level model was trained on all but one subject and evaluated on the held-out subject’s data. We quantified model fit using the average cross-validated log-likelihood across folds. State reliability across folds was evaluated via spatial correlations, with optimal state matching determined using the Hungarian algorithm (Kuhn, 1955). Pattern similarity scores were averaged across all fold pairs to yield a summary measure of cross-validated pattern reliability. We also provided the model selection result from LOOCV in Supplementary Figure 1.

HMM analyses were conducted on two experimental groups—the affair context group (n=19) and paranoia context group (n=19)—and two constructed groups—a combined group comprising all participants (n=38) and a balanced group (n=19), to match the contextual groups, consisting of random subset of participants (9 from the affair context and 10 from the paranoia context). The balanced group was created to preserve the contextual heterogeneity of the combined group while matching the sample size of the individual context groups. This approach enabled both the investigation of context-specific brain state dynamics and generalizable patterns across contexts. Brain states were characterized by spatial patterns, temporal sequence (i.e., occurrence), and inter-subject consistency. All analyses were implemented using Python (hmmlearn, NumPy, SciPy).

### State pattern similarity analysis

Traditional methods for analyzing hidden Markov models usually depend on model selection criteria to pinpoint a single optimal model, often overlooking valuable insights from alternative solutions (Quinn et al., 2018; Vidaurre, Hunt, et al., 2018). To overcome this limitation, we created a pattern similarity analysis framework that utilizes information from various model parameterizations and experimental conditions. Our approach builds on the neurobiological observation that increasing the number of states in an HMM often results in meaningful subdivisions of broader brain state processes rather than entirely spurious patterns (Baker et al., 2014). For example, a language processing state in a simpler model might subdivide into different states for different semantic domains in more complex models, representing valid phenomena at different levels of granularity.

Reliable state patterns from all HMM solutions across the experimental (affair, paranoia) and constructed groups (combined, balanced) were extracted using stringent criteria designed to filter out unstable or noisy states. First, we required a minimum activation threshold (>0.1), meaning that at least one network’s mean parameter in the HMM had to deviate from baseline by more than 0.1 (in standardized units of the z-scored input time series). This ensured that states reflected meaningful network engagement rather than near-zero fluctuations. Second, we assessed the stability of each state’s mean activation pattern using bootstrap resampling: for each network, we generated 95% confidence intervals around its mean activation value and retained only states with narrow intervals (width < 0.3), indicating low estimation uncertainty. Third, we evaluated within-state reliability using a split-half procedure: timepoints assigned to each state were divided into two halves, mean activation patterns were estimated separately, and their correlation was computed; only states with correlations > 0.5 were retained. For each retained state, we recorded fractional occupancy to standardize comparisons across models, along with full provenance information (group, model specification, original state index, normalized index).

We then clustered the state mean activation patterns across groups (Figure 2B), focusing on the average co-activation profile of the 17 networks for each state. Pairwise similarity was computed using Jaccard distance, which quantifies dissimilarity based on the proportion of non-overlapping active networks. This metric emphasizes the spatial layout of co-active brain networks while reducing sensitivity to overall activation magnitude. All states were compared pairwise, and agglomerative hierarchical clustering with average linkage was applied to group them. We evaluated clustering solutions across merging similarity thresholds from 0.6 to 0.9 and found that the top four clusters were highly stable across this range (Supplementary Figure 3).

For each consecutive pair of thresholds, we compared the consensus patterns of the top five largest clusters using Jaccard similarity (Supplementary Table 2). The analysis revealed a high degree of stability, with an average Jaccard similarity of 0.89 for the best-matching clusters across all transitions. Furthermore, 83% of these top clusters maintained a high similarity (≥ 0.7) with a cluster from the preceding threshold. This high consistency confirms that the primary state patterns are a stable feature of the data, not an artifact of parameter selection. Based on this confirmed robustness, particularly in the 0.75-0.90 range, we selected a final similarity threshold of 0.8 for all subsequent analyses.

As part of the cross-group clustering procedure, clusters were reordered by total fractional occupancy, defined as the sum of the occupancies of all states assigned to each cluster across models. This ensured that cluster IDs reflected the most frequently expressed patterns rather than simply the largest number of constituent states. The occupancy-based reordering occurred immediately after hierarchical clustering, integrated within the clustering step. For each cluster, we computed a consensus pattern identifying networks that were significantly active in at least 50% of the constituent states.

Finally, clusters were labeled as either context-general or context-specific based on their provenance. Clusters containing states from both affair and paranoia models (and appearing in the combined/balanced models) were considered context-general, whereas clusters dominated by states from only one context group were considered context-specific. To evaluate the similarity structure among brain state clusters, we computed Spearman rank correlations of their activation patterns across the 17 networks. We conducted the analysis using both consensus patterns (averaged across all member states) and representative patterns (from the combined-group models), which yielded convergent results. Here we report the correlations based on representative patterns; full results from both approaches are provided in Supplementary Figure 8.

### Story feature annotation

To examine how narrative content influenced brain state dynamics, we annotated the stimulus with key linguistic and narrative features at the temporal resolution of the fMRI data (one annotation per TR).

#### Character and interaction features

As the narrative was delivered by a single narrator but featured multiple characters, we identified character-specific speech and interactions per TR. The annotations included: (1) Arthur, Lee, and Girl speaking: Identify the intended speaking character at each time point. (2) Lee and the girl together: Identify when Lee and the girl appeared concurrently, regardless of dialogue.

#### Linguistic features

Story were tagged for grammatical parts of speech, including verbs, nouns, adjectives, and adverbs, indicating their presence in each TR.

#### Thematically relevant combined features

We further derived composite features to reflect interactions between character presence and linguistic structure: (1) Lee-Girl Verb (Lee & Girl Together × Verb Presence): Captured shared actions or relational dynamics relevant to the affair group’s expected sensitivity to relational events. (2) Arthur Adjective (Arthur Speaking × Adjective Presence): Highlighted descriptive attributes linked to Arthur, informed by findings that heightened attention to character traits is characteristic of paranoid cognition (M. J. Green & Phillips, 2004).

These structured annotations enabled the systematic evaluation of how different narrative elements influenced cognitive engagement, providing an essential foundation to investigate the hypothesized cognitive biases associated with each group.

### Bayesian generalized linear mixed models

To investigate the temporal dynamics of brain state patterns and corresponding behavioral responses, and to clarify how contextual information modulates the impact of narrative content features, we implemented Bayesian generalized linear mixed models (GLMMs). Separate GLMM analyses with identical structures were applied to characterize brain-context-content and behavior-context-content relationships, providing consistent modeling frameworks for brain and behavioral dynamics. The schematic overview of the full pipeline, from HMM to GLMM, is in Supplementary Figure 2.

### GLMM for brain state and content analysis

While the clustering analysis identified spatial configurations of brain states, the temporal dynamics necessitated identifying representative state occurrences. A representative brain state was chosen for each cluster’s first occurrence within the combined group HMMs, as these models included all participants. Subsequently, we extracted each participant’s state sequence (on/off) data corresponding to these representative states at each time point. We fit a logistic GLMM separately for each identified cluster with the following structure:

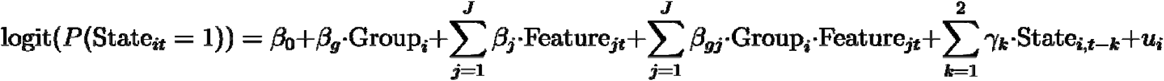

Where **State*_it_*** is a binary variable indicating whether the target brain state was active (1) or inactive (0) for subject at timepoint ***t***. **Feature*_jt_*** represents narrative annotations (e.g., character presence, linguistic elements). The interaction terms **Group*_i_*** • **Feature*_jt_*** assess whether content features affect brain state dynamics differently between groups. Autoregressive terms **State*_it-k_*** account for temporal dependencies in state occupancy, and ***u_i_*** represents subject-specific random intercepts that capture individual variability in state prevalence.

Model parameters were estimated using maximum a posteriori (MAP) estimation, applying deviation coding for group identity (+1 for affair, -1 for paranoia) and incorporating default Bayesian priors: normal priors with a mean of 0 for fixed effects and inverse gamma priors for random effects variance components. These priors provide implicit regularization, which is advantageous given our moderate sample size and binary outcomes. We calculated posterior probabilities instead of frequentist p-values for inference, quantifying evidence for effects as the probability mass supporting a specific direction of influence. This Bayesian approach allows for a more intuitive interpretation of uncertainty in our parameter estimates.

To address multiple comparisons, we implemented a Bayesian False Discovery Rate (FDR) procedure that controls the expected proportion of false discoveries among claimed discoveries. Features were considered to have credible effects when their FDR-adjusted posterior probabilities exceeded 0.95.

Coefficient estimates were converted from log-odds to odds ratios (OR) to enhance interpretability, indicating how narrative features influenced primary brain state activation odds. Group-specific effects were calculated to clarify how content features differentially affected brain state dynamics in each context condition. All analyses were performed using custom Python with the *statsmodels* package.

### GLMM for behavioral response and content analysis

We applied a generalized linear mixed model (GLMM), analogous to those used in the brain state analyses, to examine the relationship between narrative content features and behavioral responses in a separate participant sample. The dependent variable was a binary indicator reflecting whether a button press occurred at each fMRI time point (TR), signaling that the participant perceived evidence in the narrative consistent with their assigned contextual prompt. Originally recorded continuously (seconds), behavioral responses were aligned to the nearest TR to ensure temporal correspondence with stimulus features and brain-state estimates. If multiple button presses occurred within a single TR for a given participant, they were counted as a single response to avoid overrepresenting clustered inputs.

The behavioral GLMM followed this structure:

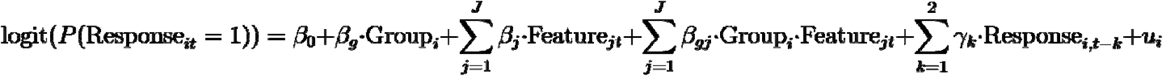

Where **Response*_it_*** is a binary variable indicating whether subject *i* pressed the button at timepoint ***t***. **Feature*_jt_*** represents narrative annotations (e.g., character presence, linguistic elements). Interaction terms **Group*_i_*** • **Feature*_jt_*** assess whether content features differentially affect behavioral responses across groups. Autoregressive terms **Response*_it-k_*** account for temporal dependencies in response patterns, and ***u_i_*** represents subject-specific random intercepts.

For parameter estimation, we utilized Maximum A Posteriori (MAP) estimation with Bayesian priors to stabilize the estimates, which is particularly important for binary outcomes with temporal dependencies. Similar to the brain-content analyses, we applied deviation coding for group identity (+1 for affair, -1 for paranoia), ensuring that the parameter estimates were balanced around the overall mean. Random intercepts at the subject level accounted for variability in individual response tendencies. Effects were deemed credible when their FDR-adjusted posterior probabilities surpassed 0.95.

## Results

### Brain state clustering identifies shared and context-specific cortical network patterns

The clustering analysis of brain state patterns across all models identified both context-specific and context-invariant brain states. Models derived from individual experimental conditions (affair or paranoia) produced more context-specific state patterns, as indicated by higher cluster IDs corresponding to smaller clusters, particularly when the number of states increased. Conversely, combined and constructed groups exhibited more generalized state patterns, represented by lower cluster IDs indicating larger clusters (Supplementary Figures 4-7). Brain states extracted from the combined group were more consistent than those extracted from the constructed balanced group, likely due to differences in sample sizes. Broad states that dominate at low model complexities (e.g., Cluster 1 in 2 to 5 state models) can fractionate into multiple, lower-occupancy sub-states at higher model complexities, which explains their apparent “disappearance” in some intermediate state solutions. This likely reflects state fractionation, a fundamental property of hierarchical brain state organization (Li et al., 2023).

Moreover, each consensus cluster represents the core defining features of each brain state pattern while accommodating natural variation within each cluster; individual states may exhibit additional network activations beyond the consensus, reflecting context-specific or model-specific nuances. For example, Cluster 2’s consensus includes auditory, control network B (Ctr-B), default mode (DMN-A/B/C), and language networks, which appear in 78–100% of its member states (Supplementary Figure 11). Some individual states within this cluster additionally recruit control network A (Ctr-A), though this occurs in only 11% of cases and thus does not define the cluster’s core identity.

The four most prominent clusters (Figure 2C), identified based on the total fractional occupancy of their constituent brain states, displayed distinct spatial configurations. Cluster 1, which had the highest total occupancy (sum=6.288), featured a representative brain state involving the Auditory, DMN-A, DMN-B, and Language networks. Cluster 2 included a representative brain state encompassing Auditory, Control-B, DMN-A, DMN-B, DMN-C, and Language networks.

Clusters 3 and 4 displayed distinct context-specific characteristics. Cluster 3 primarily comprised brain states derived from the affair context models and was largely absent in the paranoia context models. Its representative state pattern included Control-A, Control-C, Dorsal Attention-A, Dorsal Attention-B, Salience/Ventral Attention-A, Salience/Ventral Attention-B, Visual-A, and Visual-B networks. In contrast, Cluster 4 was predominantly composed of brain states from paranoia context models and was absent in the affair context models. The representative pattern of this cluster involved Control-A, Salience/Ventral Attention-A, and Salience/Ventral Attention-B networks. Details of the regions within each network can be found in Table S1.

Spearman’s rank correlations revealed positive associations between Clusters 1 and 2 (⊡ = 0.583) and between Clusters 3 and 4 (⊡ = 0.627), while Clusters 1/2 versus Clusters 3/4 were negatively correlated. Full correlation results are shown in Supplementary Figure 8.

Furthermore, Supplementary Figure 9 illustrates these relationships in low-dimensional space (PCA, MDS, t-SNE), demonstrating clear within-cluster similarity, across-cluster separation, and closer proximity between Clusters 1 and 2.

### Contextual modulation of brain state dynamics during story comprehension

Representative brain states (Figure 3) for each identified cluster were selected based on their first occurrence in the combined group HMMs: Cluster 1 (first state from 2-states model), Cluster 2 (fifth state from 7-states model), Cluster 3 (second state from 2-states model), and Cluster 4 (third state from 6-states model). These states exemplify each cluster’s core consensus pattern while potentially including additional network activations that occur in subsets of the cluster’s constituent states (Supplementary Figures 10-13). Bayesian GLMMs were then estimated separately for each cluster to determine how narrative features influenced brain state dynamics and whether these effects were modulated by context.

**Figure 3.**
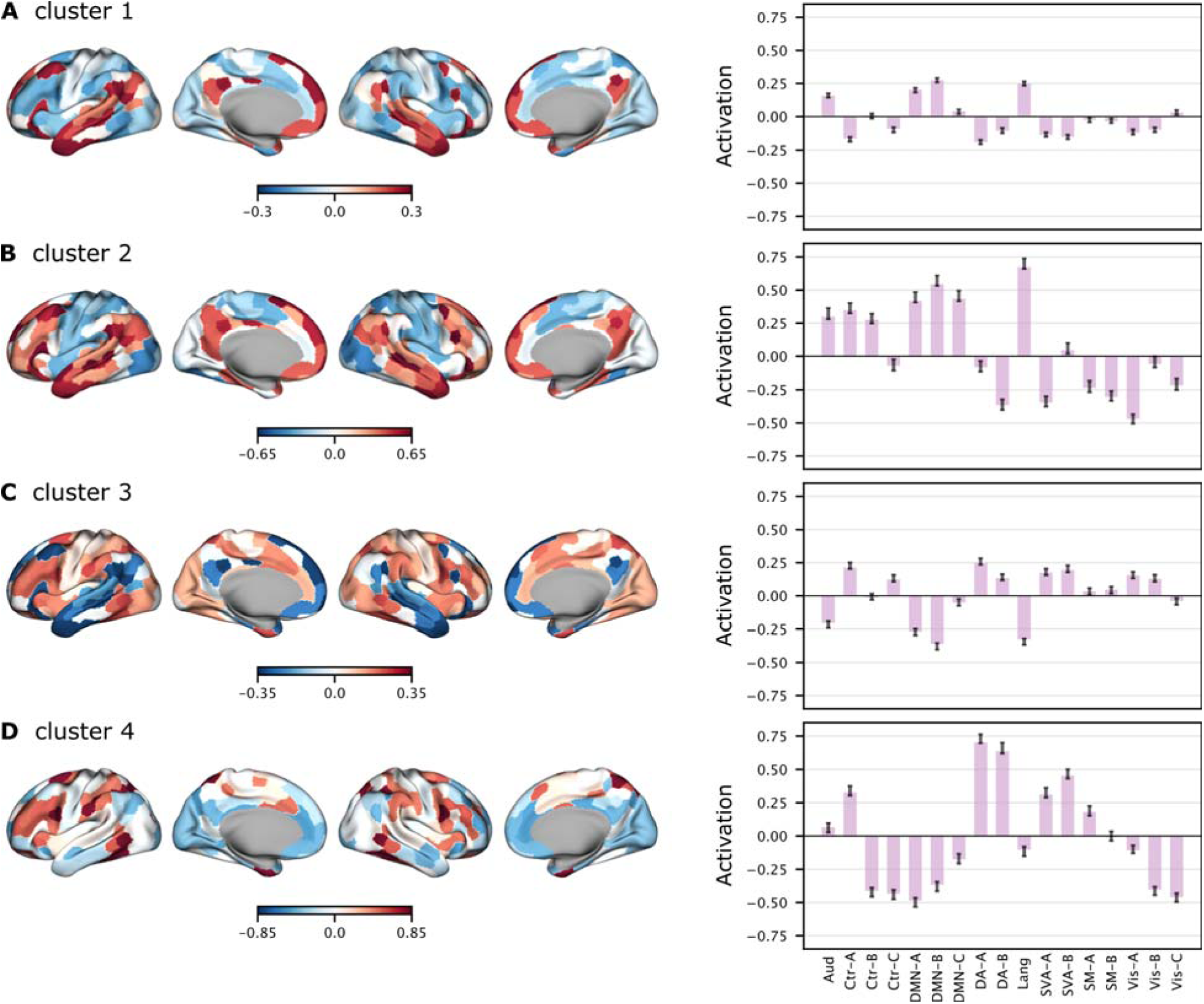
Brain activation patterns of representative brain states from the top four clusters. Each row shows a representative brain state from one of the top four clusters identified in the combined group HMMs. The left panel displays surface maps of average whole-brain activation, computed by averaging brain activity across all participants during timepoints when this state was active. Warm colors indicate above-average activation, and cool colors indicate below-average activation (z-scored). The right panel shows network-level activation profiles, summarizing average activation within canonical functional networks. Error bars represent the standard error across states belonging to the same cluster.

### Character-specific speech modulated shared brain states

Representative brain states of clusters 1 and 2 consistently presented across narrative contexts but showed different sensitivity to specific narrative features (Figure 4). In Cluster 1, Arthur speaking reliably increased the odds of Cluster 1 activation (OR=1.232, CI= [1.097, 1.383]), as did the presence of verbs (OR=1.153, CI= [1.032, 1.288]). Adjective usage showed reliable context differences (OR=0.87, CI= [0.769, 0.985]), indicating a stronger negative effect in the affair context (OR=0.833, P(Effect>0) =0.021) compared to paranoia (OR=1.1, P(Effect>0) =0.858). State occupancy was higher in the affair context (0.618) than in paranoia (0.538).

**Figure 4.**
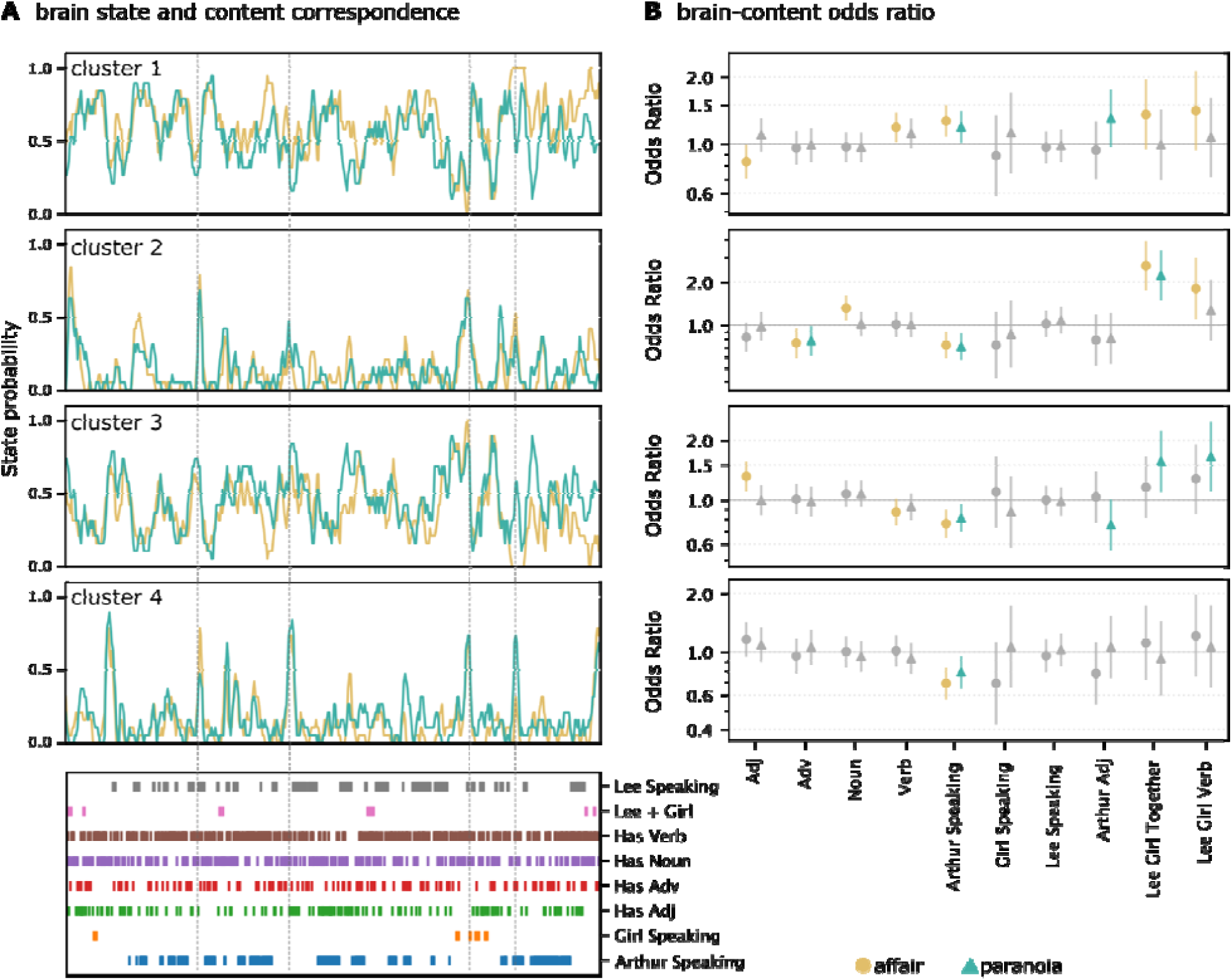
Correspondence between brain state dynamics and story content. (A) Time series of cluster-wise brain state probabilities (gold: affair group; teal: paranoia group) for the top four HMM-derived clusters in the combined group, which was estimated using all participants. Because both groups share the same latent state definitions, these traces show how strongly each state is expressed over time in the shared model, rather than providing independent group-specific estimates. Accordingly, the curves in panel A are intended to illustrate the temporal structure of each state during the narrative; statistical group differences must be evaluated in panel B rather than inferred from visual differences in these time series. The bottom panel indicates annotated linguistic and narrative features aligned to the story timeline (x-axis represent time in TR unit), including character speech and presence, part-of-speech tags (verbs, nouns, adjectives, adverbs), and key character pairings (e.g., “Lee + Girl”). (B) Bayesian logistic regression analysis estimates the odds ratios (95% credible intervals) for the association between content features and brain state cluster expression. Each panel corresponds to one of the four clusters. Points represent posterior means of the odds ratio for each content predictor; error bars show 95% credible intervals. Colored markers indicate predictors with context-dependent effects that survived FDR correction in the affair (gold) and paranoia (teal) groups. Gray markers indicate predictors without statistically reliable group differences (FDR ≥ 0.05). Odds ratios above 1 suggest an increased likelihood of brain state expression when the corresponding feature is present. Conversely, odds ratios below 1 indicate a decreased likelihood of brain state expression, meaning that the feature is associated with reduced engagement of that state.

**Figure 5.**
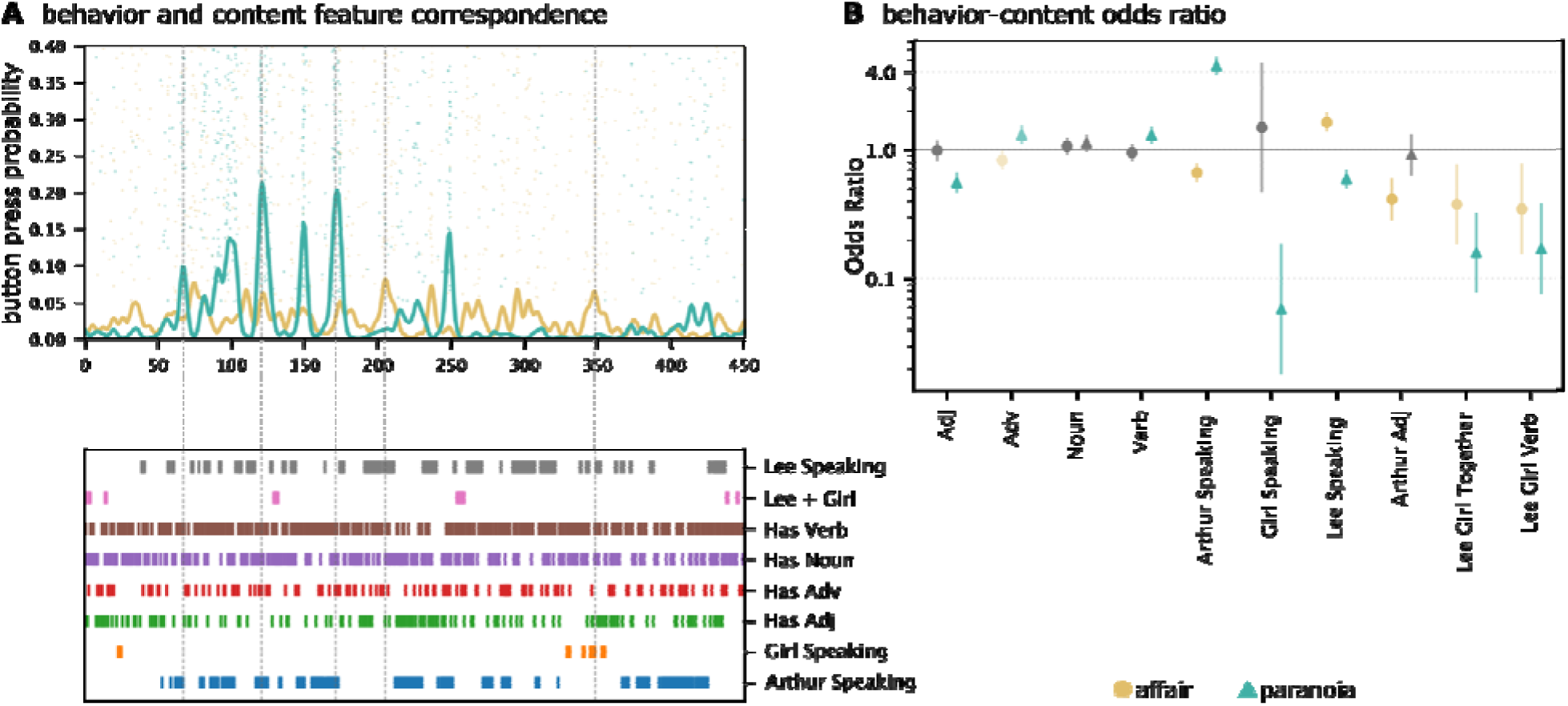
Correspondence between behavioral responses and story content features. (**A**) Time series of button press probabilities from participants in the *affair* (gold) and *paranoia* (teal) groups, reflecting the likelihood of detecting context-consistent narrative information over time. Dots represent individual button presses; smoothed lines indicate group-averaged response probabilities. The lower panel shows annotated story content features (identical to those in Figure 4A), including character speech, co-occurrence, and linguistic categories. (**B**) Bayesian logistic regression assessing the relationship between content features and button press behavior. Posterior means of odds ratios (±95% credible intervals) are plotted for each predictor. Gold and teal indicate predictors with context-dependent effects that survived FDR correction in the *affair* and paranoia groups, respectively. Gray indicates predictors without statistically reliable group differences (FDR ≥ 0.05). Odds ratios above 1 suggest an increased likelihood of brain state expression when the corresponding feature is present. Conversely, odds ratios below 1 indicate a decreased likelihood of brain state expression, meaning that the feature is associated with reduced engagement of that state.

Cluster 2 similarly showed context interactions for Arthur-related adjectives (OR=0.797, CI= [0.594, 1.069]), with odds ratios indicating a slightly stronger negative effect in the affair context (OR=0.786, P(Effect>0) =0.128) than in paranoia (OR=0.809, P(Effect>0) =0.158). Lee and girl co-present (OR=2.42, CI=[1.817, 3.221]) and their actions (OR=1.515, CI=[1.066, 2.153]) increased brain state activation odds substantially, whereas Arthur speaking (OR=0.715, CI=[0.612, 0.834]), and adverb usage (OR=0.762, CI=[0.644, 0.902]) were associated with decreases. State occupancy rate was similar in the affair context (0.121) than in paranoia (0.130).

### Context-specific brain states reveal diverse character influences

Representative brain states of clusters 3 and 4 exhibited distinct patterns that indicate context-specific processing mechanisms (Figure 4). Cluster 3 showed context-dependent modulation: Arthur speaking was associated with reduced odds of state activation (OR=0.786, CI=[0.7, 0.882]), while the combined presence (*lee_girl_together*, OR=1.348, CI=[1.045, 1.739]) of Lee and the girl with verbs (*lee_girl_verb*, OR=1.456, CI=[1.094, 1.938]) increased activation odds. Adjective usage increased the odds of Cluster 3 activation overall (OR=1.149, 95% CI [1.016, 1.3]), with odds ratios indicating a positive effect in the affair context (OR=1.316, P(Effect>0) =0.999) but negligible effects in paranoia (OR=0.997, P(Effect>0) =0.486). State occupancy was lower in the affair context (0.382) than paranoia (0.462).

Cluster 4 showed context interactions for Arthur speaking (OR=0.936, CI= [0.814, 1.076]). Arthur speaking demonstrated a slightly stronger negative effect in the affair context (OR=0.692, P(Effect>0) =0) compared to paranoia (OR=0.79, P(Effect>0) =0.009). Overall state occupancy was slightly lower in the affair context (0.137) than paranoia (0.171).

Multiple comparison analyses with Bayesian False Discovery Rate (FDR) corrections confirmed that the odds ratios for adjective usage (Clusters 1 and 3), Arthur-related adjectives (Clusters 1 and 2), and Arthur speaking (Cluster 4) consistently differed from chance expectations (FDR < 0.05). These results provide credible evidence that narrative context strongly modulates the temporal dynamics of brain states, revealing distinct narrative feature influences across different contexts.

### Contextual modulation of behavioral responses during story comprehension

Bayesian GLMMs showed that odds ratios for participants’ button presses differed systematically across contexts, indicating contextual modulation of behavioral responses.

### Context-dependent impact of character identity on behavioral responses

Character-related narrative features showed strong overall influences on behavioral responses, with Arthur speaking (OR=2.743, CI= [2.286, 3.291]) and Lee speaking (OR=1.801, CI= [1.508, 2.150]) substantially increasing the probability of button presses. Critically, context strongly modulated these effects. Arthur speaking showed large context-dependent differences in odds (interaction OR=0.334, CI= [0.279, 0.401]), with substantially greater response likelihood in the paranoia context (OR=8.205, P(Effect>0) =1.000) compared to minimal effects in the affair context (OR=0.917, P(Effect>0) =0.254). Similarly, Girl speaking was associated with higher odds in the affair context but lower odds in the paranoia context (interaction OR=2.864, CI=[1.189, 6.894]), positively influencing responses in the affair context (OR=1.559, P(Effect>0)=0.758) but strongly reducing responses in the paranoia context (OR=0.190, P(Effect>0)=0.004). Lee speaking also showed credible contextual modulation (interaction OR=0.813, CI= [0.681, 0.970]), with a stronger effect in the paranoia context (OR=2.216, P(Effect>0)>0.999) relative to the affair context (OR=1.463, P(Effect>0) =0.999).

### Context-specific effects of linguistic features

Linguistic features displayed smaller credible context-dependent patterns in predicting behavioral responses. Noun usage increased the odds of button presses overall (OR=1.351, CI= [1.205, 1.515]), modulated by context (interaction OR=0.811, CI= [0.723, 0.909]). Nouns more strongly increased response probability in the paranoia context (OR=1.667, P(Effect>0)>0.999) than in the affair context (OR=1.095, P(Effect>0) =0.865). Similarly, adverb usage showed context-dependent modulation of odds (interaction OR=0.814, CI= [0.717, 0.924]), negatively influencing button presses in the affair context (OR=0.844, P(Effect>0) =0.032) but positively in paranoia (OR=1.275, P(Effect>0) =0.996). Verb usage had a modest positive main effect (OR=1.046, CI=[0.933, 1.172]) and credible context interaction (interaction OR=0.917, CI=[0.818, 1.027]), with stronger effects observed in paranoia (OR=1.141, P(Effect>0)=0.946) than in the affair context (OR=0.959, P(Effect>0)=0.303).

### Descriptive character features show contextual variation

Arthur-related adjectives were associated with reduced odds of responses overall (OR=0.411, CI= [0.297, 0.570]), with credible contextual modulation (interaction OR=1.155, CI= [0.834, 1.599]). The negative effect was stronger in the affair context (OR=0.475, P(Effect>0) =0.001) compared to paranoia (OR=0.356, P(Effect>0) <0.001). The combined presence of Lee and girl characters negatively influenced responses (OR=0.445, CI= [0.168, 1.175]), though this context modulation did not meet FDR criteria. The interaction of these characters with verbs showed no credible context-specific effects. Supplementary Materials 2.3 confirms that the behavioral GLMM results replicate at 1-s resolution.

## Discussion

Our study investigated how narrative context modulates brain state dynamics and behavioral responses during story comprehension. We identified both shared and context-specific brain states that spanned auditory, language, default mode, control, attention, and visual networks. By modeling the relationships between stimulus features, context, and brain states, we found credible evidence that context influences how narrative features, particularly speech or references to specific characters, impact the temporal dynamics of these brain states. Independent behavioral analyses revealed context-dependent differences in odds ratios for stimulus features, indicating selective modulation of behavioral responses by context.

### Shared brain states suggest convergent processing during story listening

Clusters 1 and 2 reflected DMN-dominant narrative-integration states that appeared consistently across both context conditions, in line with our prediction that naturalistic story comprehension would elicit stable integration-focused brain states. Both clusters showed strong involvement of auditory and language networks, which support the processing of the unfolding speech signal, as well as DMN-A and DMN-B regions, including the medial prefrontal cortex (PFCm), posterior cingulate cortex (PCC), inferior parietal lobule (IPL), precuneus (pCun), and temporal and temporopolar cortices. These DMN subsystems have been widely implicated in long-timescale semantic integration, situation-model construction, and narrative coherence (Hasson et al., 2018; Jackson et al., 2023; Raichle et al., 2001; Simony et al., 2016). Together, these profiles suggest that Clusters 1 and 2 may capture a core processing mode through which incoming linguistic information is incrementally incorporated into a coherent representation of the evolving narrative.

Cluster 2 slightly differed from Cluster 1 by additionally recruiting regions within control network-B, such as dorsolateral prefrontal cortex, inferior frontal gyrus, intraparietal sulcus, and orbitofrontal cortex, as well as posterior DMN-C regions including the retrosplenial/parahippocampal cortex and posterior precuneus. Control network-B is associated with adaptive cognitive control, strategic monitoring, and attentional modulation (Cole et al., 2014; Dworetsky et al., 2024), while DMN-C has been linked to contextual updating, mental simulation, and scene construction (Ritchey & Cooper, 2020). These additional contributions suggest that Cluster 2 may reflect a more demanding substate of narrative integration in which listeners must reorganize their situation model, integrate newly informative plot elements, or resolve referential ambiguity. The stronger activation amplitudes observed in Cluster 2 are therefore consistent with theories proposing that the DMN comprises multiple specialized subsystems that support both continuous integration (DMN-A/B) and event-level updating or simulation (DMN-C), and that transitions into control-network engagement mark moments of locally increased cognitive demand. In this view, Cluster 1 represents a dominant “baseline” integration mode, whereas Cluster 2 reflects brief but intensive episodes of narrative updating supported by coordinated engagement of control and higher-order DMN subsystems.

### Context-specific brain states index specialized processing demands

Clusters 3 and 4 reflected MDN-dominant brain states that were differentially expressed across the two context conditions. This pattern is consistent with longstanding accounts of the multiple-demand system (MDN), which emphasize that interpretive effort dynamically recruits distinct subcomponents of control, dorsal attention, and salience networks depending on task demands (Duncan, 2010; Cole et al., 2013; Fedorenko et al., 2013; Uddin, 2015). Unlike the DMN-based integration states (Clusters 1 and 2), these MDN states showed minimal involvement of auditory, language, or core DMN regions, perhaps reflecting evaluative, attentionally demanding, and context-sensitive modes of narrative processing.

Cluster 3, expressed predominantly in the affair group, engaged regions within control network-C, including the posterior cingulate cortex, precuneus, lateral prefrontal cortex, medial prefrontal cortex, and parahippocampal cortex, and precuneus, together with visual association areas in Visual Networks A and B. Although mean activation levels in this state were relatively low, the joint involvement of posterior midline control regions and visual cortices may implicate a visual suggests a simulation-oriented or imagery-based mode of narrative evaluation in an ambiguous narrative (Koide-Majima et al., 2024; Liu et al., 2022; Pearson, 2019). Cluster 4, in contrast, appeared primarily in the paranoia group and exhibited a strong, canonical MDN profile, with pronounced activation in control, dorsal attention, and salience networks (Duncan, 2010; Cole et al., 2013). These networks support executive vigilance, uncertainty monitoring, and the detection of behaviorally relevant or potentially threatening cues (Hermans et al., 2014; Uddin, 2015).

The correlation structure among clusters suggests that these four states can be understood along a single functional axis ranging from context-general to context-sensitive modes of processing. This pattern aligns with prior work showing that low-dimensional decompositions of brain activity reveal recurrent state families that organize around stable, large-scale network configurations (Bolt et al., 2022; Song et al., 2023). In our case, Clusters 1 and 2 correspond to DMN-dominant integration states that appeared across both groups, supporting continuous processes such as semantic integration, situation-model construction, and the incorporation of incoming narrative information. In contrast, Clusters 3 and 4 represent distinct MDN-related evaluative states whose expression diverged across the primed contexts. These findings support our hypothesis that naturalistic story comprehension involves both stable DMN-supported integration states and context-sensitive MDN-supported evaluative states, with the latter varying systematically according to the interpretive demands elicited by the priming manipulation.

### Narrative context modulates the influence of story features on brain state dynamics

Our Bayesian GLMM analyses revealed that several narrative features showed credible, feature-specific differences in their associations with brain state activation across the two context conditions (Figure 4). Across all clusters, state probabilities were reliably modulated by narrative content, consistent with prior work demonstrating that linguistic and narrative cues exert moment-to-moment influences on neural processing during story comprehension (Jacoby & Fedorenko, 2020; Yarkoni et al., 2008). However, the specific features exerting these effects, and the magnitude and direction of those effects, varied across clusters and contexts.

For Clusters 1 and 2, which appeared across both narrative contexts, we observed modest but credible feature-specific interactions. In Cluster 1, Arthur speaking and verb usage increased activation probability, whereas adjective usage showed a negative context interaction, indicating a stronger decrease in the affair condition than in paranoia. Cluster 2 showed similar scattered interactions: Arthur-related adjectives exhibited weak context differences, while features related to the co-presence and actions of Lee and the girl strongly increased activation, and adverbs and Arthur speaking reduced it. These effects suggest that the DMN-dominant integration states remain broadly engaged across contexts but show subtle variations in their sensitivity to specific linguistic cues, consistent with accounts proposing that even stable integrative processes are modulated by moment-to-moment narrative content (Grall & Finn, 2022; Mar, 2011).

Clusters 3 and 4 exhibited somewhat clearer context-dependent patterns, with several content features influencing state activation differently across the two groups. In Cluster 3, Arthur speaking reduced activation odds, while co-occurrence and action features related to Lee and the girl increased activation, and adjectives showed a positive effect primarily in the affair condition. Cluster 4 showed credible context differences for Arthur speaking, with a stronger negative effect in the affair group. These findings indicate that, for these clusters, context influences how specific narrative cues shape the likelihood of entering a given state. However, the effects remain modest in magnitude, typical for naturalistic fMRI datasets, and do not reflect global shifts in state identity but rather differences in how particular narrative features drive transient shifts into states with distinct functional profiles.

To assess whether these context-dependent patterns in the representative states could arise from arbitrary inter-individual variability, we conducted permutation analyses comparing observed group differences against a null distribution derived from 10,000 random participant splits (Supplementary Figure 14). For Clusters 1, 3, and 4, observed differences in temporal dynamics exceeded 95% of random splits (all p < .05), confirming that context-based grouping produces systematically different state engagement than would be expected from chance. Cluster 2 did not show this effect (p = .249). Cluster 2 shares core DMN and language network involvement with Cluster 1 but additionally engages Control-B and DMN-C regions. One possibility is that this configuration supports narrative-tracking processes, such as updating situational details or monitoring discourse coherence, that proceed similarly regardless of the listener’s interpretive frame. However, we cannot rule out that the finer model granularity from which the representive Cluster 2 state derives (from the combined-group 7-state HMM) contributes to reduced sensitivity for detecting group differences.

Taken together, these findings suggest a two-level architecture for context effects in narrative comprehension. At the level of state engagement, context biases which network configurations are preferentially recruited over time with Clusters 1, 3, and 4 showing temporal dynamics that systematically differ by interpretive frame. At the level of feature sensitivity, both groups show largely similar responses to narrative content, with selective divergence for features requiring evaluative inference (particularly adjectives). This pattern indicates that context does not dramatically or continuously restructure how the brain processes narrative content over time. Rather, context operates as a modulatory bias: it shapes the probability of engaging particular network configurations while much of the underlying content-processing architecture is preserved. The features that do show context-dependent modulation, such as descriptive language conveying character traits and evaluative information, are those for which prior expectations should matter most, as their interpretation requires integration with the listener’s model of what happened or what the story “is about.”

### Contextual modulation of behavioral responses highlights selective engagement with narrative content

Independent behavioral analyses reinforce our observation that narrative context selectively modulates which stimulus features influence responses. For example, odds ratios indicated that Arthur’s speaking substantially increased the likelihood of button presses in the paranoia context, but not in the affair context. This sensitivity to character speech aligns with established evidence from narrative psychology and psycholinguistics, showing heightened audience engagement with central narrative figures who guide interpretive frameworks and ensure story coherence (Eekhof et al., 2023; M. C. Green & Appel, 2024; Hartung et al., 2017). Likewise, Lee’s speaking elicited strong context-dependent behavioral responses, reflecting variations in perceived narrative relevance or emotional significance of different characters across contexts.

Importantly, these character-driven effects in behavioral responses were stronger than those observed in brain-state analyses, suggesting distinct cognitive mechanisms underlying explicit versus implicit narrative processing. Explicit behavioral responses likely reflect deliberate inferential and evaluative processes, such as active narrative coherence assessments, conscious attribution of narrative significance, or explicit inference generation. These explicit processes may differ from the processes evoked by passive story listening.

Linguistic features such as nouns and adverbs exhibited modest but reliable context-dependent effects in behavioral analyses. These differences indicate that certain lexical categories may be differentially weighted or processed depending on the narrative context (Tilmatine et al., 2024), though the specific mechanisms, whether semantic, evaluative, or otherwise, remain to be clarified. Descriptive adjectives showed more substantial negative effects in the affair context. This pattern is consistent with prior psycholinguistic findings suggesting that adjectives can modulate emotional tone and guide interpretive inferences, particularly in context-sensitive narratives (Lei et al., 2023).

These findings demonstrate that prior contextual information systematically modulated behavioral responses to narrative content, with the strongest effects observed for speech attribution and more modest effects for specific linguistic features. The pattern suggests that context does not uniformly enhance or suppress engagement with all content features but instead modulates behavioral responses in a selective, feature-dependent manner.

A final consideration concerns the apparent misalignment between the fMRI and behavioral findings: whereas the behavioral analyses showed strong and directionally consistent context effects of linguistic and character features, the fMRI-derived brain state dynamics did not closely follow these behavioral directions, even though they showed statistically reliable modulation. This difference reflects the fact that the two paradigms, while based on the same narrative stimulus, are fundamentally different tasks. The fMRI experiment involved passive listening, followed by delayed comprehension questions, which captured implicit, distributed interpretive processes without discrete response markers. By contrast, the behavioral experiment required explicit evidence detection with overt button presses, producing stronger and more directional effects as participants actively relied on linguistic cues for decision-making. Because the tasks differ in both cognitive demands and data type (neural dynamics vs. behavioral reports), a one-to-one alignment of results is not expected. The behavioral task would be expected to engage decision- and motor-related systems not central to naturalistic comprehension. Taken together, the behavioral findings extend the fMRI results by showing how linguistic features guide explicit reasoning, while the brain-state analyses reveal implicit neural dynamics of narrative comprehension.

### Limitations and future directions

Our study provides valuable insights into how narrative context modulates brain state dynamics and behavioral responses, yet several limitations point toward promising avenues for future research. First, our HMMs were conducted at the network level, averaging signals within the 17 predefined functional networks. This approach is well-motivated, given that large-scale networks are reliable units of naturalistic fMRI analysis, but it necessarily reduces spatial resolution. Finer-grained analyses (e.g., ROI-level or voxelwise HMMs; Vidaurre, Abeysuriya, et al., 2018) could reveal more heterogeneous dynamics within networks and provide stronger leverage for testing hypotheses about default mode (DMN) and multiple-demand (MDN) subsystem involvement; note that the 17 pre-defined functional networks used here were derived from resting-state parcellation and thus reflect intrinsic functional organization rather than task-evoked subdivisions. At the same time, higher spatial granularity substantially increases model dimensionality and can lead to unstable HMM estimation for naturalistic datasets of this length (451 TRs). Thus, the network-level approach represents a pragmatic and theoretically grounded compromise, with higher-resolution approaches remaining an important target for future work.

Another limitation is that standard HMMs assume a geometric distribution of dwell times, which tends to bias results toward shorter state durations and more frequent switching between states. Hidden semi-Markov models (HSMMs) address this limitation by modeling dwell times explicitly, offering more accurate estimates of state persistence (Shappell et al., 2019). Future work could also employ hierarchical nonparametric variants such as the hierarchical Dirichlet process HMM (HDP-HMM), which infers both the number of states and their duration properties directly from the data (Beal et al., 2002; Fox et al., 2011). These approaches would provide a more flexible framework for examining naturalistic brain dynamics beyond the assumptions of conventional HMMs.

A further methodological limitation concerns model initialization. In this study, state means were initialized with random draws from a normal distribution rather than with more structured approaches such as k-means clustering, which can sometimes improve stability for high state counts. While our validation procedures (cross-validation, bootstrap uncertainty estimation, clustering across models) minimize the risk that results depend on any single initialization scheme, future work could compare alternative initialization strategies to assess their impact on reproducibility and model fit.

In addition, aspects of our stimulus (e.g., its suspenseful narrative structure) and experimental design (e.g., contextual priming) introduce features that are unique to this study. Although the two broad categories of brain-state clusters we identified, DMN- and MDN-related, are likely to generalize across similar naturalistic story-listening paradigms, the finer-grained configuration of these states (e.g., relative activation amplitudes, involvement of sensory networks, or the emergence of sub-states) may vary across studies. Such variability likely reflects differences in narrative content, stimulus structure, and experimental design. Future work would benefit from a large-scale mega-analysis across existing story-listening fMRI datasets to characterize how DMN- and MDN-related brain states vary as a function of stimulus features and study design.

Moreover, our neuroimaging and behavioral data were collected from separate participant samples. Although both datasets independently revealed context-sensitive effects, collecting brain and behavioral responses from the same individuals would allow for tighter linkage between neural state dynamics and subjective narrative judgments, enabling more direct tests of brain–behavior relationships (Xu et al., 2025). We also note that the dependent variables in the brain and behavioral GLMMs differ: brain models use a binary indicator of state activation per subject at each timepoint, whereas behavioral models reflect the proportion of participants in each group who pressed a key at each moment. Because the scales and noise properties of these measures differ, brain–behavior comparisons should be interpreted as convergent rather than one-to-one correspondences. Furthermore, the binary nature of the brain dependent variable may limit sensitivity to detect graded context-dependent modulations; continuous measures of state expression could reveal subtler effects that discrete classification obscures. Future studies could address this by acquiring brain and behavioral data in the same individuals or by using hierarchical joint modeling with continuous state metrics.

While we used part-of-speech (PoS) labels to characterize contextual differences in language input, these labels offer only a shallow approximation of meaning. PoS categories reflect syntactic structure rather than semantic content, and future work should incorporate richer linguistic features, such as word embeddings, semantic role labels, or discourse structure, to better capture the narrative elements that drive brain dynamics.

Finally, using a single narrative stimulus may limit the generalizability of our findings to other narrative forms or genres. Future studies examining diverse narrative types, varying in emotional content, complexity, and modality, could test the breadth and boundaries of context effects on brain dynamics and behavioral engagement.

Together, these extensions would refine our understanding of how context shapes neural and behavioral responses to narrative and support more general models of naturalistic cognition.

## Conclusion

This study provides converging neural and behavioral evidence that contextual framing shapes how listeners process and interpret unfolding narrative information. Using brain-state modeling, we identified recurrent states that were expressed across both groups and engaged auditory, language, and default mode networks consistent with ongoing narrative integration, alongside additional states whose temporal expression varied with contextual priming and showed differentiated sensitivity to specific narrative features. These context-related differences were modest in magnitude, as is typical for naturalistic fMRI data, but statistically credible and aligned with feature-dependent distinctions observed in the behavioral task. The behavioral findings similarly showed that character-related cues, particularly direct speech, influenced participants’ interpretive judgments in a context-dependent manner. Together, these results suggest that narrative context shapes comprehension not only retrospectively but also through subtle, moment-to-moment adjustments in how linguistic features influence large-scale brain dynamics. By integrating state-based neural modeling with time-resolved behavioral measures, this work provides an initial foundation for understanding how contextual framing interacts with narrative structure to guide ongoing cognitive processing.

## Data and Code Availability

The Python code used for our analysis and visualization is available at https://github.com/yibeichan/prettymouth

## Author Contributions

YC: Conceptualization, Methodology, Software, Formal analysis, Investigation, Data Curation, Visualization, Writing - Original Draft, Project administration. ZZ: Conceptualization, Visualization, Resources, Writing - Review & Editing. SN: Conceptualization, Resources, Writing - Review & Editing, Supervision. GA: Conceptualization, Writing - Review & Editing, Supervision. SG: Funding acquisition, Resources, Writing - Review & Editing, Supervision.

## Funding

YC and SSG were partially supported by NIH P4 EB019936 and by the Lann and Chris Woehrle Psychiatric Fund at the McGovern Institute for Brain Research at MIT.

## Acknowledgements

We would like to thank William Menegas and the members of the Senseable Intelligence Group for insightful discussions.

## Declaration of Competing Interest

The authors declare that they have no known competing financial interests or personal relationships that could have appeared to influence the work reported in this paper.

## Notes

### Competing Interest Statement

The authors have declared no competing interest.

### Summary of Updates

In this revision, we made significant updates to the Introduction and Discussion to clarify our theoretical framework and provide more precise interpretations of our findings. In particular, we (i) articulated a clearer DMN- and MDN-related rationale for expecting distinct classes of brain states during narrative processing in the Introduction; (ii) revised the Discussion to interpret state-level differences more carefully and GLMM results without overstating their magnitude; and (iii) integrated a new summary of how context shapes feature-state relationships in a content-specific rather than global manner. We also improved clarity around the HMM and GLMM analyses. We explicitly described how representative state time series were derived from the combined-group HMM, clarified the role of the content-feature x context interaction in identifying context effects, and revised figure captions (especially for Figure 4) to avoid potential misinterpretation.

