## supplementary_material for "Context modulates brain state dynamics and behavioral responses during narrative comprehension"

### 1. Methods

#### 1.1 fMRI data

We utilized the “prettymouth” dataset, consisting of 40 participants drawn from the Narrative dataset (Nastase et al., 2021). Participants were divided into two groups (initially N = 20 per group), each exposed to an adapted version of J. D. Salinger's short story, "Pretty Mouth and Green My Eyes." The adapted version was shorter than the original and included several sentences not present in the original text. Narration was provided by a professional actor, resulting in a recording lasting 11 minutes and 32 seconds. Functional MRI data were acquired with a repetition time (TR) of 1.5 seconds. The story was preceded by 18 seconds of neutral music and 3 seconds of silence, followed by an additional 15 seconds. These segments of music and silence were excluded from all analyses.

The narrative describes a phone conversation between two friends, Arthur and Lee. Arthur, who has just returned home from a party after losing track of his wife Joanie, calls Lee to express his concerns about her whereabouts. Lee is at home with a woman beside him, whose identity remains ambiguous—she may or may not be Joanie. Before listening to the story, each participant group received different contextual information: one group was informed that Arthur was paranoid and his suspicions were unfounded (paranoia context), while the other group was told that the woman was indeed Joanie, Arthur’s wife, and that Lee and Joanie had been involved in an ongoing affair for over a year (affair context). Yeshurun et al. (2017) and Nastase et al. (2021) describe the experimental paradigm and fMRI data acquisition parameters.

#### 1.2 fMRI data preprocessing

Results included in this manuscript come from preprocessing performed using *fMRIPrep* 24.1.0 (Esteban et al. (2019); Esteban et al. (2018); RRID:SCR_016216), which is based on *Nipype* 1.8.6 (K. Gorgolewski et al. (2011); K. J. Gorgolewski et al. (2018); RRID:SCR_002502).

##### 1.2.1 Anatomical data preprocessing

A total of 2 T1-weighted (T1w) images were found within the input BIDS dataset. Each T1w image was corrected for intensity non-uniformity (INU) with N4BiasFieldCorrection (Tustison et al. 2010), distributed with ANTs 2.5.3 (Avants et al. 2008, RRID:SCR_004757). The T1w-reference was then skull-stripped with a *Nipype* implementation of the antsBrainExtraction.sh workflow (from ANTs), using OASIS30ANTs as target template. Brain tissue segmentation of cerebrospinal fluid (CSF), white-matter (WM) and gray-matter (GM) was performed on the brain-extracted T1w using fast (FSL (version unknown), RRID:SCR_002823, Zhang, Brady, and Smith 2001). An anatomical T1w-reference map was computed after registration of 2 <module 'nipype.interfaces.image' from '/opt/conda/envs/fmriprep/lib/python3.11/site-packages/nipype/interfaces/image.py'> images (after INU-correction) using mri_robust_template (FreeSurfer 7.3.2, Reuter, Rosas, and Fischl 2010). Brain surfaces were reconstructed using recon-all (FreeSurfer 7.3.2, RRID:SCR_001847, Dale, Fischl, and Sereno 1999), and the brain mask estimated previously was refined with a custom variation of the method to reconcile ANTs-derived and FreeSurfer-derived segmentations of the cortical gray-matter of Mindboggle (RRID:SCR_002438, Klein et al. 2017). Volume-based spatial normalization to two standard spaces (MNI152NLin6Asym, MNI152NLin2009cAsym) was performed through nonlinear registration with antsRegistration (ANTs 2.5.3), using brain-extracted versions of both T1w reference and the T1w template. The following templates were were selected for spatial normalization and accessed with *TemplateFlow* (24.2.0, Ciric et al. 2022): *FSL's MNI ICBM 152 non-linear 6th Generation Asymmetric Average Brain Stereotaxic Registration Model* [Evans et al. (2012), RRID:SCR_002823; TemplateFlow ID: MNI152NLin6Asym], *ICBM 152 Nonlinear Asymmetrical template version 2009c* [Fonov et al. (2009), RRID:SCR_008796; TemplateFlow ID: MNI152NLin2009cAsym]. *Grayordinate* "dscalar" files containing 91k samples were resampled onto fsLR using the Connectome Workbench (Glasser et al. 2013).

**1.2.2 Preprocessing of boinhomogeneity mappings**

A total of 3 fieldmaps were found available within the input BIDS structure for this particular subject. A deformation field to correct for susceptibility distortions was estimated based on *fMRIPrep*'s *fieldmap-less* approach. The deformation field is that resulting from co-registering the EPI reference to the same-subject T1w-reference with its intensity inverted (Wang et al. 2017; Huntenburg 2014). Registration is performed with antsRegistration (ANTs 2.5.3), and the process regularized by constraining deformation to be nonzero only along the phase-encoding direction, and modulated with an average fieldmap template (Treiber et al. 2016).

**1.2.3 Functional data preprocessing**

For each of the 3 BOLD runs found per subject (across all tasks and sessions), the following preprocessing was performed. First, a reference volume was generated, using a custom methodology of *fMRIPrep*, for use in head motion correction. Head-motion parameters with respect to the BOLD reference (transformation matrices, and six corresponding rotation and translation parameters) are estimated before any spatiotemporal filtering using mcflirt (FSL , Jenkinson et al. 2002). The estimated *fieldmap* was then aligned with rigid-registration to the target EPI (echo-planar imaging) reference run. The field coefficients were mapped on to the reference EPI using the transform. The BOLD reference was then co-registered to the T1w reference using bbregister (FreeSurfer) which implements boundary-based registration (Greve and Fischl 2009). Co-registration was configured with six degrees of freedom. Several confounding time-series were calculated based on the *preprocessed BOLD*: framewise displacement (FD), DVARS and three region-wise global signals. FD was computed using two formulations following Power (absolute sum of relative motions, Power et al. (2014)) and Jenkinson (relative root mean square displacement between affines, Jenkinson et al. (2002)). FD and DVARS are calculated for each functional run, both using their implementations in *Nipype* (following the definitions by Power et al. 2014). The three global signals are extracted within the CSF, the WM, and the whole-brain masks. Additionally, a set of physiological regressors were extracted to allow for component-based noise correction (*CompCor*, Behzadi et al. 2007). Principal components are estimated after high-pass filtering the *preprocessed BOLD* time-series (using a discrete cosine filter with 128s cut-off) for the two *CompCor* variants: temporal (tCompCor) and anatomical (aCompCor). tCompCor components are then calculated from the top 2% variable voxels within the brain mask. For aCompCor, three probabilistic masks (CSF, WM and combined CSF+WM) are generated in anatomical space. The implementation differs from that of Behzadi et al. in that instead of eroding the masks by 2 pixels on BOLD space, a mask of pixels that likely contain a volume fraction of GM is subtracted from the aCompCor masks. This mask is obtained by dilating a GM mask extracted from the FreeSurfer's *aseg* segmentation, and it ensures components are not extracted from voxels containing a minimal fraction of GM. Finally, these masks are resampled into BOLD space and binarized by thresholding at 0.99 (as in the original implementation). Components are also calculated separately within the WM and CSF masks. For each CompCor decomposition, the *k* components with the largest singular values are retained, such that the retained components' time series are sufficient to explain 50 percent of variance across the nuisance mask (CSF, WM, combined, or temporal). The remaining components are dropped from consideration. The head-motion estimates calculated in the correction step were also placed within the corresponding confounds file. The confound time series derived from head motion estimates and global signals were expanded with the inclusion of temporal derivatives and quadratic terms for each (Satterthwaite et al. 2013). Frames that exceeded a threshold of 0.9 mm FD or 5.0 standardized DVARS were annotated as motion outliers. Additional nuisance timeseries are calculated by means of principal components analysis of the signal found within a thin band (*crown*) of voxels around the edge of the brain, as proposed by (Patriat, Reynolds, and Birn 2017). The BOLD time-series were resampled onto the left/right-symmetric template "fsLR" using the Connectome Workbench (Glasser et al. 2013). *Grayordinates* files (Glasser et al. 2013) containing 91k samples were also generated with surface data transformed directly to fsLR space and subcortical data transformed to 2 mm resolution MNI152NLin6Asym space. All resamplings can be performed with *a single interpolation step* by composing all the pertinent transformations (i.e. head-motion transform matrices, susceptibility distortion correction when available, and co-registrations to anatomical and output spaces). Gridded (volumetric) resamplings were performed using nitransforms, configured with cubic B-spline interpolation.

Many internal operations of *fMRIPrep* use *Nilearn* 0.10.4 (Abraham et al. 2014, RRID:SCR_001362), mostly within the functional processing workflow. For more pipeline details, see the section corresponding to workflows in *fMRIPrep*'s documentation.

**1.2.4 Functional data post-processing**

After fMRIPrep (version 24.1.0), we conducted custom post-processing optimized for narrative listening analyses. Specifically, we applied spatial smoothing (6 mm FWHM) to balance noise reduction and preservation of spatial activation patterns, detrended the data to remove scanner drift, standardized (z-scored) signals within-subject, and removed nuisance signals associated with motion and physiological noise using motion parameters and aCompCor regressors. Post-processing was conducted using nilearn (version 0.10.1), detailed code can be found here (https://github.com/yibeichan/prettymouth/blob/main/script/01_postproc.py). Data extraction was performed using Schaefer2018_1000Parcels_Kong2022_17Networks parcellation (https://github.com/ThomasYeoLab/CBIG/blob/master/stable_projects/brain_parcellation/Schaef er2018_LocalGlobal/Parcellations/MNI/Schaefer2018_1000Parcels_Kong2022_17Networks_order_FSLMNI152_2mm.nii.gz).

**Supplementary Table 1. 17 networks and names of parcels in those networks**

| **Networks** | **Parcel names** |
|---|---|
| Auditory | Ins (Insula), ParOcc (Parietal Occipital), ParOper (Parietal Operculum), ST (Superior Temporal), Temp (Temporal), TempPole (Temporal Pole) |
| ContA (control network A) | Cingm (Mid-Cingulate), IPS (Intraparietal Sulcus), Ins (Insula), OFC (Orbital Frontal Cortex), PFCd (Dorsal Prefrontal Cortex), PFCl (Lateral Prefrontal Cortex), PFCmp (Medial Posterior Prefrontal Cortex), SPL (Superior Parietal Lobule), Temp (Temporal) |
| ContB (control network B) | FPole (Frontal Pole), IFG (Inferior Frontal Gyrus), IPL (Inferior Parietal Lobule), IPS (Intraparietal Sulcus), OFC (Orbital Frontal Cortex), PCC (Posterior Cingulate Cortex), PFCd (Dorsal Prefrontal Cortex), PFCl (Lateral Prefrontal Cortex), PFCmp (Medial Posterior Prefrontal Cortex), Temp (Temporal) |
| ContC (control network C) | Cingp (Cingulate Posterior), FPole (Frontal Pole), IPL (Inferior Parietal Lobule), Ins (Insula), OFC (Orbital Frontal Cortex), PCC (Posterior Cingulate Cortex), PFCl (Lateral Prefrontal Cortex), PFCm (Medial Prefrontal Cortex), PHC (Parahippocampal Cortex), pCun (Precuneus) |
| DefaultA (default mode network A) | FPole (Frontal Pole), IPL (Inferior Parietal Lobule), PCC (Posterior Cingulate Cortex), PFCd (Dorsal Prefrontal Cortex), PFCm (Medial Prefrontal Cortex), PHC (Parahippocampal Cortex), Temp (Temporal), TempPole (Temporal Pole), pCun (Precuneus) |
| DefaultB (default mode network B) | FPole (Frontal Pole), IPL (Inferior Parietal Lobule), PCC (Posterior Cingulate Cortex), PFCd (Dorsal Prefrontal Cortex), PFCl (Lateral Prefrontal Cortex), PFCv (Ventral Prefrontal Cortex), Temp |

| | |
|---|---|
| | (Temporal), TempPole (Temporal Pole), pCun (Precuneus) |
| DefaultC (default mode network C) | IPL (Inferior Parietal Lobule), PFCd (Dorsal Prefrontal Cortex), PHC (Parahippocampal Cortex), RSC (Retrosplenial Cortex), pCun (Precuneus) |
| DorsAttnA (dorsal attention network A) | IPL (Inferior Parietal Lobule), IPS (Intraparietal Sulcus), PFCd (Dorsal Prefrontal Cortex), PrCd (Precentral Dorsal), PrCv (Precentral Ventral), SPL (Superior Parietal Lobule), TempOcc (Temporal Occipital), TempPole (Temporal Pole) |
| DorsAttnB (dorsal attention network B) | FrMed (Frontal Medial), IPL (Inferior Parietal Lobule), ParOper (Parietal Operculum), PostC (Post Central), PrCd (Precentral Dorsal), PrCv (Precentral Ventral), SPL (Superior Parietal Lobule) |
| Language | IFG (Inferior Frontal Gyrus), IPL (Inferior Parietal Lobule), PFCd (Dorsal Prefrontal Cortex), PFCl (Lateral Prefrontal Cortex), Temp (Temporal), TempPole (Temporal Pole) |
| SalVenAttnA (salience ventral attention network A) | FrMed (Frontal Medial), FrOper (Frontal Operculum), IPL (Inferior Parietal Lobule), Ins (Insula), PFCd (Dorsal Prefrontal Cortex), PFCmp (Medial Posterior Prefrontal Cortex), SPL (Superior Parietal Lobule) |
| SalVenAttnB (salience ventral attention network B) | IFG (Inferior Frontal Gyrus), IPL (Inferior Parietal Lobule), Ins (Insula), OFC (Orbital Frontal Cortex), PFCd (Dorsal Prefrontal Cortex), PFCl (Lateral Prefrontal Cortex), PFCmp (Medial Posterior Prefrontal Cortex), SPL (Superior Parietal Lobule), Temp (Temporal) |
| SomMotA (Somatosensory Motor network A) | ParOper (Parietal Operculum) |
| SomMotB (Somatosensory | Ins (Insula), ParOper (Parietal Operculum) |

| | |
|---|---|
| Motor network B) | |
| VisualA | ExStr (Extrastriate Cortex), PrC (Precentral), SPL (Superior Parietal Lobule) |
| VisualB | ExStrInf (Extra-striate Inferior), ExStrSup (Extra-striate Superior), SPL (Superior Parietal Lobule), Striate (Striate Cortex) |
| VisualC | ExStr (Extrastriate Cortex), Striate (Striate Cortex) |

##### 1.3 Behavioral data collection

Behavioral data were collected from 128 participants recruited via Prolific (www.prolific.co). Participants were classified into two experimental groups: an affair group (n = 63) and a paranoia group (n = 59). Demographic details indicated the sample comprised 62 males, 57 females, and three individuals identifying as non-binary or third gender. The age distribution included participants aged 18–24 years (n = 18), 25–34 years (n = 46), 35–44 years (n = 29), 45–54 years (n = 12), 55–64 years (n = 14), and 65 years or older (n = 3).

Data were collected through an online experiment developed using PsychoJS scripts derived from the PsychoPy builder (PsychoPy3, version 2023.2.0) and hosted on Pavlovia (https://pavlovia.org/). Participants initially provided informed consent via Qualtrics (https://www.qualtrics.com/) before being randomly assigned to one of two context conditions (affair versus paranoia). Participants then listened to an audio story while receiving instructions identical to those during the neuroimaging experiment.

**Supplementary Textbox 1. Instructions on Pavlovia**

Following a 5-second countdown, you will hear a brief musical interlude, after which you will listen to the following story:

Affair version:

It is late at night and the phone is ringing. On one end of the line is Arthur; Arthur just came home from a party. He left the party without finding his wife, Joanie. As always, Joanie was flirting with everybody at the party. Arthur is very upset. On the other end is Lee, Arthur's friend. He is at home with Joanie, Arthur's wife. Lee and Joanie have just returned from the same party. They have been having an affair for over a year now. They are thinking about the excuse Lee will use to calm Arthur this time.

Paranoia version:

It is late at night and the phone is ringing. On one end of the line is Arthur; Arthur just came home from a party. He left the party without finding his wife, Joanie. As always, Arthur is paranoid, worrying that she might be having an affair, which is not true. On the other end is Lee, Arthur's friend. He is at home with his girlfriend, Rose. Lee and Rose have just returned from the same party, and are desperate to go to sleep. They do not know anything about Joanie's whereabouts, and are tired of dealing with Arthur's overreactions.

Your task is to *press the SPACE bar on your keyboard whenever you perceive information that suggests Lee and Joanie are having an affair (Arthur is being paranoid) (based on the description of the situation above)*. When you respond to confirm your input, a green dot will appear on your screen.

Once you finish the above task, you will be redirected to the second part of the experiment: a questionnaire where you will be asked to answer questions about the story.

When you are ready to begin, please press 'Enter/Return' on your keyboard to initiate the task.

This study was approved by the Princeton University Institutional Review Board (IRB 12201). All participants provided informed consent electronically before participation, in accordance with institutional ethical guidelines. Participants received monetary compensation consistent with university policy. All data were anonymized to ensure participant confidentiality.

##### 1.4 Hidden Markov model analysis

We employed Hidden Markov Models (HMMs) to examine brain state dynamics during narrative comprehension. HMMs are particularly well-suited for capturing state transitions in continuous cognitive processes, as they identify recurring spatial patterns of brain activity without imposing rigid temporal constraints (Vidaurre et al., 2017; Baldassano et al., 2018). To ensure the robustness and generalizability of the identified states, we implemented cross-validation procedures and statistical assessments across participants.

###### 1.4.1 Data preprocessing

Functional MRI data underwent standard preprocessing, including motion correction, spatial normalization, and spatial smoothing. Time series data were extracted from network-defined parcels, yielding a multidimensional representation of brain activity for each participant (n=19 per group) across 475 timepoints. To account for hemodynamic delays, the BOLD time series were temporally shifted forward by three TRs (~4.5 s) to better align brain activity with the recorded signal (Yeshurun et al., 2017). Additionally, non-story segments—consisting of background music or silence spanning 24 TRs at both the beginning and end of each scan—were excluded from analysis. This preprocessing step resulted in 451 TRs available for subsequent HMM input. To mitigate individual differences in BOLD signal amplitude, each participant's time series was independently z-scored prior to HMM analysis, ensuring that state identification was based on relative activation patterns rather than absolute signal magnitudes.

**1.4.2 HMM framework**

We implemented a Gaussian observation HMM in which each brain state was modeled as generating multivariate Gaussian-distributed activity patterns with state-specific means and covariances. The modeling framework was developed using the hmmlearn Python package, incorporating methodological refinements to enhance reliability and mitigate potential biases in state estimation.

Model parameters were initialized as follows. State means were drawn from a standard normal distribution $\mathcal{N}(0, 1)$ for each state and feature, ensuring broad coverage of the feature space. Covariance matrices were initialized as identity matrices with a small diagonal ridge term to guarantee positive definiteness and numerical stability. Start probabilities were initialized uniformly. Transition probabilities were initialized to favor an expected dwell time of ~7 s (~4–5 TRs), using the formula $p_{ii} = \exp(-1/d)$ where d is the expected duration in TRs, with remaining probability mass distributed uniformly across off-diagonal transitions. This provided a weak prior consistent with evidence that brain states during naturalistic cognition typically persist on the order of several seconds, with higher-order regions exhibiting longer durations and sensory regions switching more rapidly (Vidaurre et al., 2017; Baldassano et al., 2017). Transition probabilities were subsequently re-estimated during expectation–maximization, so final dwell times were determined by the data.

To reduce sensitivity to local optima, we ran five independent optimization attempts with different random seeds, which affected both the random initialization of state means and the EM algorithm's internal stochastic path. For each state number, we selected the solution with the highest log-likelihood.

##### 1.4.3 Cross-validation procedure

To evaluate model generalizability and determine the optimal number of states, we conducted leave-one-subject-out cross-validation (LOOCV). In each iteration, the model was trained on all but one participant and evaluated on the held-out subject. Model performance was assessed by quantifying the consistency of identified state patterns across cross-validation folds. Spatial correlation analyses were performed to evaluate state similarity, with the Hungarian algorithm used to address label permutation issues. Log-likelihood scores on held-out data provided an additional measure of predictive performance. To further assess state reliability, we examined the reproducibility of identified states across training partitions, ensuring that they reflected stable brain activity patterns rather than subject-specific artifacts.

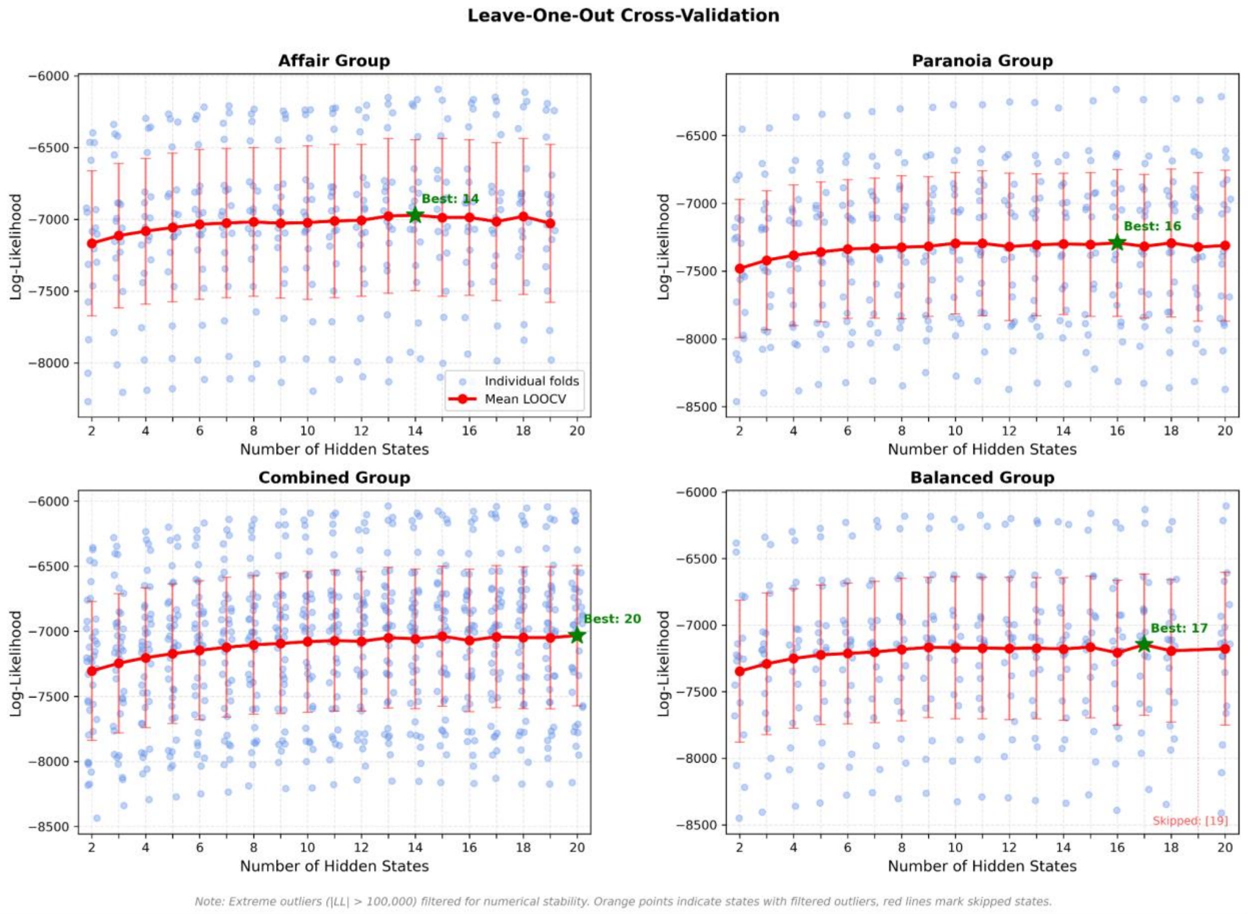


**Supplementary Figure 1. Model selection via leave-one-out cross-validation.**
Each panel shows LOOCV results for one group: (a) Affair (n = 19), (b) Paranoia (n = 19), (c) Combined (n = 38), and (d) Balanced (n = 38, subsampled). Blue points are individual fold log-likelihoods, and the red line (±1 SD) shows the group mean; gold stars mark the state number

with the highest mean log-likelihood. Outlier folds due to numerical instabilities are shown in orange and were excluded. Optimal state numbers were 14, 16, 20, and 17 for the Affair, Paranoia, Combined, and Balanced groups, though 6–12 states showed comparable performance with greater interpretability. The Balanced 19-state model failed to converge, and the Affair group lacks a 20-state solution.

##### 1.4.4 State characterization

We characterized brain states for each model regarding their spatial properties, temporal dynamics, and inter-subject consistency. Spatial analyses included examination of state activation patterns, the reliability of these patterns across cross-validation folds, and the separability of states in high-dimensional space using Euclidean and Mahalanobis distance metrics. Temporal properties were assessed by computing state durations, transition probabilities, switching rates, and recurrence intervals. Additionally, we evaluated the degree to which state occurrences and sequences were shared across participants, quantifying inter-subject synchronization in state dynamics.

##### 1.4.5 Uncertainty estimation

Statistical uncertainty was estimated using bootstrap resampling with 1,000 iterations at the participant level. This non-parametric approach provided confidence intervals for state-related metrics while accounting for inter-individual variability without assuming a specific distribution for the underlying parameters.

HMM analyses were performed on four groups: the affair context group (n=19), the paranoia context group (n=19), a combined group that included all participants from both contexts (n=38), and a constructed balanced group (n=19) formed by randomly selecting 9 subjects from the affair context and 10 from the paranoia context. This constructed group allowed for balanced representation while controlling for sample size effects when comparing against individual context groups. This approach allowed us to examine context-specific brain state dynamics and general patterns shared across contexts. Brain states were characterized by spatial configuration, temporal properties (durations, transition probabilities, switching rates), and inter-subject consistency. All analyses were implemented in Python (hmmlearn, NumPy, SciPy).

#### 1.5 State pattern similarity analysis

Traditional approaches to hidden Markov model (HMM) analysis typically rely on model selection based on criteria such as Bayesian Information Criterion (BIC) or cross-validated likelihood, ultimately choosing a single model with a fixed number of states (Vidaurre et al., 2018; Quinn et al., 2018). However, this strategy discards potentially valuable insights contained within alternative model solutions. Neural activation patterns that consistently emerge across multiple model parameterizations might represent robust, biologically meaningful states, even if they occur in models not globally optimal according to typical selection metrics (Ryali et al., 2016; Taghia et al., 2018).

Our method leverages this observation by systematically examining brain states across various HMM parameterizations. Neurobiologically, increasing the number of states in an HMM typically results in subdividing broader cognitive states into more granular, meaningful subdivisions rather than generating entirely spurious states (Baker et al., 2014; Vidaurre et al., 2017). For example, a language-processing state identified in a simpler model could subdivide into separate syntactic and semantic processing states in more complex models, each reflecting valid neural phenomena at distinct levels of detail. Analyzing these patterns across models with varying state numbers provides stronger evidence of their neurobiological validity than single-model approaches.

**1.5.1 Pattern extraction and validation**

For two experimental groups (affair, paranoia) and two constructed groups (combined and balanced), we extracted state patterns from all available HMM solutions across different model parameterizations, irrespective of overall model-fit criteria. Each extracted pattern represented a spatial configuration of brain activity across network-defined parcels, corresponding to a specific state in an HMM solution. To ensure the reliability of identified states, we applied multiple stringent criteria:

1. Activation threshold: A state pattern required a maximum absolute activation ≥ 0.1.
2. Confidence interval width: The maximum width of the bootstrap confidence intervals across activated features was limited to ≤0.3.

3. Pattern stability: Stability was assessed using a split-half correlation method. State data were split into two halves, and the correlation between mean activation patterns from each half was computed. Stability scores ranged from 0 to 1, with a minimum required threshold of 0.5.

States with higher stability ($\geq 0.8$) were permitted slightly more lenient confidence intervals due to their demonstrated reproducibility across model iterations. Significant state features were determined based on consistent activation exceeding the threshold and robust statistical evidence from bootstrap confidence intervals. For highly stable patterns (stability $> 0.8$), activation above the threshold was considered sufficient, while less stable patterns required both threshold activation and statistical reliability, as evidenced by positive lower confidence bounds.

**1.5.2 State standardization and provenance tracking**

For each identified state pattern, we maintained comprehensive provenance information, including:

1. Originating experimental group (affair, paranoia, or combined)
2. Model specification (including the number of states)
3. Original state index within the model
4. Normalized state index (after sorting by fractional occupancy)
5. Fractional occupancy value
6. Pattern stability score
7. Active features and their activation values

To facilitate meaningful comparisons across models with different parameterizations, states within each model were sorted by fractional occupancy, with the most frequently occurring states assigned the lowest indices. This standardization procedure was critical for addressing a common challenge in cross-model state comparison (Ryali et al., 2016), ensuring that comparisons were not confounded by arbitrary state numbering.

The sorting process involved:

1. Extracting fractional occupancy values for each state from group-level metrics
2. Creating pairs of (state_idx, occupancy) for each state
3. Sorting these pairs by occupancy in descending order
4. Creating a mapping from original state indices to sorted indices

5. Storing both the mapping and occupancy information for each model

This approach ensures that states with similar functional roles (as indicated by their temporal prevalence) are compared across models, regardless of their arbitrary initial numbering.

##### 1.5.3 Cross-group pattern clustering

After extracting and standardizing state patterns from all models and groups, we performed hierarchical clustering to identify similar patterns across experimental conditions. We employed the Jaccard distance metric, which quantifies dissimilarity based on the proportion of non-shared active features between patterns:

$$\text{Jaccard distance} = 1 - |A \cap B| / |A \cup B|$$

Where A and B represent the sets of active features in two patterns, this metric was chosen for its sensitivity to differences in the topological configuration of activated brain regions, providing a more neurobiologically interpretable assessment by focusing explicitly on activation-pattern overlap and reducing sensitivity to model-specific scaling factors. The clustering procedure consisted of:

1. Computing the Jaccard distance matrix between all pairs of patterns using SciPy's pdist function
2. Performing hierarchical clustering using average linkage (UPGMA) with SciPy's linkage function
3. Cutting the dendrogram at a distance threshold of 0.3 (corresponding to a similarity threshold of 0.7)
4. We performed this clustering analysis with various thresholds, including 0.6, 0.65, 0.7, 0.75, 0.8, 0.85, and 0.9. The results of the top 5 clusters remained consistent across different thresholds.
5. To ensure that our results were not dependent on a specific hyperparameter, we conducted a stability analysis across a range of clustering thresholds (0.6 to 0.9 similarity). For each consecutive pair of thresholds, we compared the consensus patterns of the top five largest clusters using Jaccard similarity (**Supplementary Table 2**). The analysis revealed a high degree of stability, with an average Jaccard similarity of 0.89 for the best-matching clusters across all transitions. Furthermore, 83% of these top clusters maintained a high similarity (≥ 0.7) with a cluster from the preceding threshold.

6. This high consistency confirms that the primary state patterns are a stable feature of the data, not an artifact of parameter selection. Based on this confirmed robustness, particularly in the 0.75-0.90 range, we selected a final similarity threshold of 0.8 for all subsequent analyses reported in this manuscript.

**Supplementary Table 2. Cluster stability across thresholds.**

| threshold_from | threshold_to | rank | size | n_networks | networks | best_match_similarity | prev_rank |
|---|---|---|---|---|---|---|---|
| 0.6 | 0.65 | 1 | 51 | 6 | Aud, Ctr-B, DMN-A, DMN-B, DMN-C, Lang | 0.8333333333333330 | 1 |
| 0.6 | 0.65 | 2 | 47 | 8 | Ctr-A, Ctr-C, DA-A, SVA-A, SVA-B, Vis-A, Vis-B, Vis-C | 1.0 | 2 |
| 0.6 | 0.65 | 3 | 39 | 5 | Ctr-A, DA-A, DA-B, SVA-A, SVA-B | 1.0 | 3 |
| 0.6 | 0.65 | 4 | 37 | 4 | Aud, DMN-A, DMN-B, Lang | 0.8 | 1 |
| 0.6 | 0.65 | 5 | 32 | 15 | Aud, Ctr-B, Ctr-C, DMN-A, DMN-B, DMN-C, DA-A, DA-B, Lang, SVA-A, SM-A, SM-B, Vis-A, Vis-B, Vis-C | 0.4117647058823530 | 5 |
| 0.65 | 0.7 | 1 | 39 | 5 | Ctr-A, DA-A, DA-B, SVA-A, SVA-B | 1.0 | 3 |
| 0.65 | 0.7 | 2 | 37 | 4 | Aud, DMN-A, DMN-B, Lang | 1.0 | 4 |
| 0.65 | 0.7 | 3 | 36 | 6 | Aud, Ctr-B, DMN-A, DMN-B, DMN-C, Lang | 1.0 | 1 |
| 0.65 | 0.7 | 4 | 23 | 6 | Aud, DA-B, SVA-A, SVA-B, SM-A, SM-B | 0.375 | 3 |
| 0.65 | 0.7 | 5 | 23 | 8 | Aud, Ctr-A, Ctr-B, Ctr-C, DMN-A, DMN-B, DMN-C, Lang | 0.75 | 1 |
| 0.7 | 0.75 | 1 | 33 | 4 | Aud, DMN-A, DMN-B, Lang | 1.0 | 2 |
| 0.7 | 0.75 | 2 | 31 | 5 | Ctr-A, DA-A, DA-B, SVA-A, SVA-B | 1.0 | 1 |
| 0.7 | 0.75 | 3 | 29 | 6 | Aud, Ctr-B, DMN-A, DMN-B, DMN-C, Lang | 1.0 | 3 |
| 0.7 | 0.75 | 4 | 22 | 9 | Ctr-A, Ctr-B, Ctr-C, DA-A, SVA-A, SVA-B, Vis-A, Vis-B, Vis-C | 0.4 | 1 |
| 0.7 | 0.75 | 5 | 19 | 5 | Ctr-A, Ctr-B, Ctr-C, DMN-A, DMN-C | 0.625 | 5 |
| 0.75 | 0.8 | 1 | 24 | 4 | Aud, DMN-A, DMN-B, Lang | 1.0 | 1 |

| | | | | | | | |
|---|---|---|---|---|---|---|---|
| 0.75 | 0.8 | 2 | 24 | 5 | Ctr-A, DA-A, DA-B, SVA-A, SVA-B | 1.0 | 2 |
| 0.75 | 0.8 | 3 | 18 | 6 | Aud, Ctr-B, DMN-A, DMN-B, DMN-C, Lang | 1.0 | 3 |
| 0.75 | 0.8 | 4 | 15 | 5 | Ctr-A, Ctr-B, Ctr-C, DMN-A, DMN-C | 1.0 | 5 |
| 0.75 | 0.8 | 5 | 14 | 8 | Ctr-A, Ctr-C, DA-A, SVA-A, SVA-B, Vis-A, Vis-B, Vis-C | 0.88888888 88888890 | 4 |
| 0.8 | 0.85 | 1 | 24 | 4 | Aud, DMN-A, DMN-B, Lang | 1.0 | 1 |
| 0.8 | 0.85 | 2 | 18 | 5 | Ctr-A, DA-A, DA-B, SVA-A, SVA-B | 1.0 | 2 |
| 0.8 | 0.85 | 3 | 15 | 6 | Aud, Ctr-B, DMN-A, DMN-B, DMN-C, Lang | 1.0 | 3 |
| 0.8 | 0.85 | 4 | 15 | 5 | Ctr-A, Ctr-B, Ctr-C, DMN-A, DMN-C | 1.0 | 4 |
| 0.8 | 0.85 | 5 | 10 | 4 | DA-A, Vis-A, Vis-B, Vis-C | 0.5 | 5 |
| 0.85 | 0.9 | 1 | 24 | 4 | Aud, DMN-A, DMN-B, Lang | 1.0 | 1 |
| 0.85 | 0.9 | 2 | 18 | 5 | Ctr-A, DA-A, DA-B, SVA-A, SVA-B | 1.0 | 2 |
| 0.85 | 0.9 | 3 | 15 | 5 | Ctr-A, Ctr-B, Ctr-C, DMN-A, DMN-C | 1.0 | 4 |
| 0.85 | 0.9 | 4 | 13 | 6 | Aud, Ctr-B, DMN-A, DMN-B, DMN-C, Lang | 1.0 | 3 |
| 0.85 | 0.9 | 5 | 10 | 4 | DA-A, Vis-A, Vis-B, Vis-C | 1.0 | 5 |

##### 1.5.4 Cluster characterization and consensus pattern derivation

After hierarchical clustering, clusters were reordered by total fractional occupancy, defined as the sum of the occupancies of all member states across models. This occupancy-based reordering was integrated directly into the clustering step rather than performed afterward, and cluster IDs were reassigned accordingly. This ensured that clusters were labeled in a way that prioritized patterns most frequently expressed across models, rather than simply those with the largest number of constituent states.

For each cluster, we then derived a consensus pattern representing the common set of active features shared among member states. This consensus was computed by averaging the binary activation maps across all member patterns and thresholding at 0.5, thereby identifying features active in most contributing states. Clusters were characterized according to:

1. Prominence (total fractional occupancy): the aggregate occupancy of all member states.
2. Size: the number of constituent patterns.
3. Group composition: the distribution of states originating from each experimental group.
4. Consensus pattern: the shared set of active features.
5. Member provenance: detailed metadata linking back to each constituent state.

This procedure preserved complete bidirectional mapping between clusters and their member states, enabling us to examine how different experimental groups contributed to each cluster, to track how increasing model complexity subdivided states across clusters, and to assess which brain networks were most consistently involved in each cluster.

To quantitatively examine relationships among brain state clusters, we performed Spearman rank correlation analyses on cluster-averaged activation patterns. Two complementary approaches were employed to ensure robustness.

(1) Consensus pattern approach. For each cluster, all member state patterns (activation vectors across 17 networks) were aggregated, and the mean pattern was computed. This provided a robust representation of the cluster's characteristic activation profile by leveraging the diversity of states assigned to each cluster.

(2) Representative pattern approach. A single representative state was selected from the combined-group HMM solutions for each cluster (Cluster 1: 2-state model, state 0; Cluster 2: 7-state model, state 4; Cluster 3: 2-state model, state 1; Cluster 4: 6-state model, state 3). This provided well-defined exemplars of each cluster's activation profile.

For both approaches, Spearman rank correlations were computed between all pairs of cluster patterns, represented as 17-dimensional vectors of mean activation across canonical brain networks. Based on visual inspection, we tested three specific hypotheses: (1) positive correlation between Clusters 1 and 2, (2) positive correlation between Clusters 3 and 4, and (3) negative correlation between the two groups ([C1+C2]/2 vs. [C3+C4]/2). Statistical significance was assessed using two-tailed tests at an $\alpha$ level of 0.05.

The analysis used several Python libraries, including NumPy for numerical operations, SciPy for distance calculations, hierarchical clustering, and correlation calculation, Pandas for data manipulation, and Matplotlib and Seaborn for visualization. All code was optimized for computational efficiency, with a particular focus on memory management when handling large datasets.

#### 1.6 Story feature annotation

To examine how narrative content influenced brain state dynamics, we annotated the stimulus with key linguistic and narrative features at the temporal resolution of the fMRI data (one annotation per TR).

**Character and interaction features:** As the narrative was delivered by a single narrator but featured multiple characters, we identified character-specific speech and interactions per TR. The annotations included: (1) Arthur, Lee, and Girl speaking: Identify the intended speaking character at each time point. (2) Lee and the girl together: Identify when Lee and the girl appeared concurrently, regardless of dialogue.

**Linguistic features:** Story were tagged for grammatical parts of speech, including verbs, nouns, adjectives, and adverbs, indicating their presence in each TR.

**Thematically relevant combined features:** We further derived composite features to reflect interactions between character presence and linguistic structure: (1) Lee-Girl Verb (Lee & Girl Together × Verb Presence): Captured shared actions or relational dynamics relevant to the affair group's expected sensitivity to relational events. (2) Arthur Adjective (Arthur Speaking × Adjective Presence): Highlighted descriptive attributes linked to Arthur, informed by findings that heightened attention to character traits is characteristic of paranoid cognition (M. J. Green & Phillips, 2004).

These structured annotations enabled the systematic evaluation of how different narrative elements influenced cognitive engagement, providing an essential foundation to investigate the hypothesized cognitive biases associated with each group.

#### 1.7 Bayesian generalized linear mixed models

To investigate the temporal dynamics of brain state patterns and corresponding behavioral responses, and to clarify how contextual information modulates the impact of narrative content features, we implemented Bayesian generalized linear mixed models (GLMMs). Separate GLMM analyses with identical structures were applied to characterize brain-context-content and behavior-context-content relationships, providing consistent modeling frameworks for brain and behavioral dynamics.

##### 1.7.1 GLMM for brain state and content analysis

While the clustering analysis identified spatial configurations of brain states, the temporal dynamics necessitated identifying representative state occurrences. A representative brain state was chosen for each cluster's first occurrence within the combined group HMMs, as these models included all participants. Subsequently, we extracted each participant's state sequence (on/off) data corresponding to these representative states at each time point. We fit a logistic GLMM separately for each identified cluster with the following structure:

$$\text{logit}(P(\text{State}_{it} = 1)) = \beta_0 + \beta_g \cdot \text{Group}_i + \sum_{j=1}^{J} \beta_j \cdot \text{Feature}_{jt} + \sum_{j=1}^{J} \beta_{gj} \cdot \text{Group}_i \cdot \text{Feature}_{jt} + \sum_{k=1}^{2} \gamma_k \cdot \text{State}_{i,t-k} + u_i$$

Where $\text{State}_{it}$ is a binary variable indicating whether the target brain state was active (1) or inactive (0) for subject $i$ at timepoint $t$. $\text{Feature}_{jt}$ represents narrative annotations (e.g., character presence, linguistic elements). The interaction terms $\text{Group}_i \cdot \text{Feature}_{jt}$ assess whether content features affect brain state dynamics differently between groups. Autoregressive terms $\text{State}_{i,t-k}$ account for temporal dependencies in state occupancy, and $u_i$ represents subject-specific random intercepts that capture individual variability in state prevalence.

Model parameters were estimated using maximum a posteriori (MAP) estimation, applying deviation coding for group identity (+1 for affair, -1 for paranoia) and incorporating default Bayesian priors: normal priors with a mean of 0 for fixed effects and inverse gamma priors for random effects variance components. These priors provide implicit regularization, which is advantageous given our moderate sample size and binary outcomes. We calculated posterior probabilities instead of frequentist p-values for inference, quantifying evidence for effects as the probability mass supporting a specific direction of influence. This Bayesian approach allows for a more intuitive interpretation of uncertainty in our parameter estimates.

To address multiple comparisons, we implemented a Bayesian False Discovery Rate (FDR) procedure that controls the expected proportion of false discoveries among claimed discoveries. Features were considered to have credible effects when their FDR-adjusted posterior probabilities exceeded 0.95.

Coefficient estimates were converted from log-odds to odds ratios (OR) to enhance interpretability, indicating how narrative features influenced primary brain state activation odds. Group-specific effects were calculated to clarify how content features differentially affected brain state dynamics in each context condition. All analyses were performed using custom Python with the *statsmodels* package.

##### 1.7.2 GLMM for behavioral response and content analysis

We applied a generalized linear mixed model (GLMM), analogous to those used in the brain state analyses, to examine the relationship between narrative content features and behavioral responses in a separate participant sample. The dependent variable was a binary indicator reflecting whether a button press occurred at each fMRI time point (TR), signaling that the participant perceived evidence in the narrative consistent with their assigned contextual prompt. Originally recorded continuously (seconds), behavioral responses were aligned to the nearest TR to ensure temporal correspondence with stimulus features and brain-state estimates. If multiple button presses occurred within a single TR for a given participant, they were counted as a single response to avoid overrepresenting clustered inputs.

The behavioral GLMM followed this structure:

$$\text{logit}(P(\text{Response}_{it} = 1)) = \beta_0 + \beta_g \cdot \text{Group}_i + \sum_{j=1}^{J} \beta_j \cdot \text{Feature}_{jt} + \sum_{j=1}^{J} \beta_{gj} \cdot \text{Group}_i \cdot \text{Feature}_{jt} + \sum_{k=1}^{2} \gamma_k \cdot \text{Response}_{i,t-k} + u_i$$

Where $\text{Response}_{it}$ is a binary variable indicating whether subject $i$ pressed the button at timepoint $t$. $\text{Feature}_{jt}$ represents narrative annotations (e.g., character presence, linguistic elements). Interaction terms $\text{Group}_i \cdot \text{Feature}_{jt}$ assess whether content features differentially affect behavioral responses across groups. Autoregressive terms $\text{Response}_{i,t-k}$ account for temporal dependencies in response patterns, and $u_i$ represents subject-specific random intercepts.

For parameter estimation, we utilized Maximum A Posteriori (MAP) estimation with Bayesian priors to stabilize the estimates, which is particularly important for binary outcomes with temporal dependencies. Similar to the brain-content analyses, we applied deviation coding for group identity (+1 for affair, -1 for paranoia), ensuring that the parameter estimates were balanced around the overall mean. Random intercepts at the subject level accounted for variability in individual response tendencies. Effects were deemed credible when their FDR-adjusted posterior probabilities surpassed 0.95.

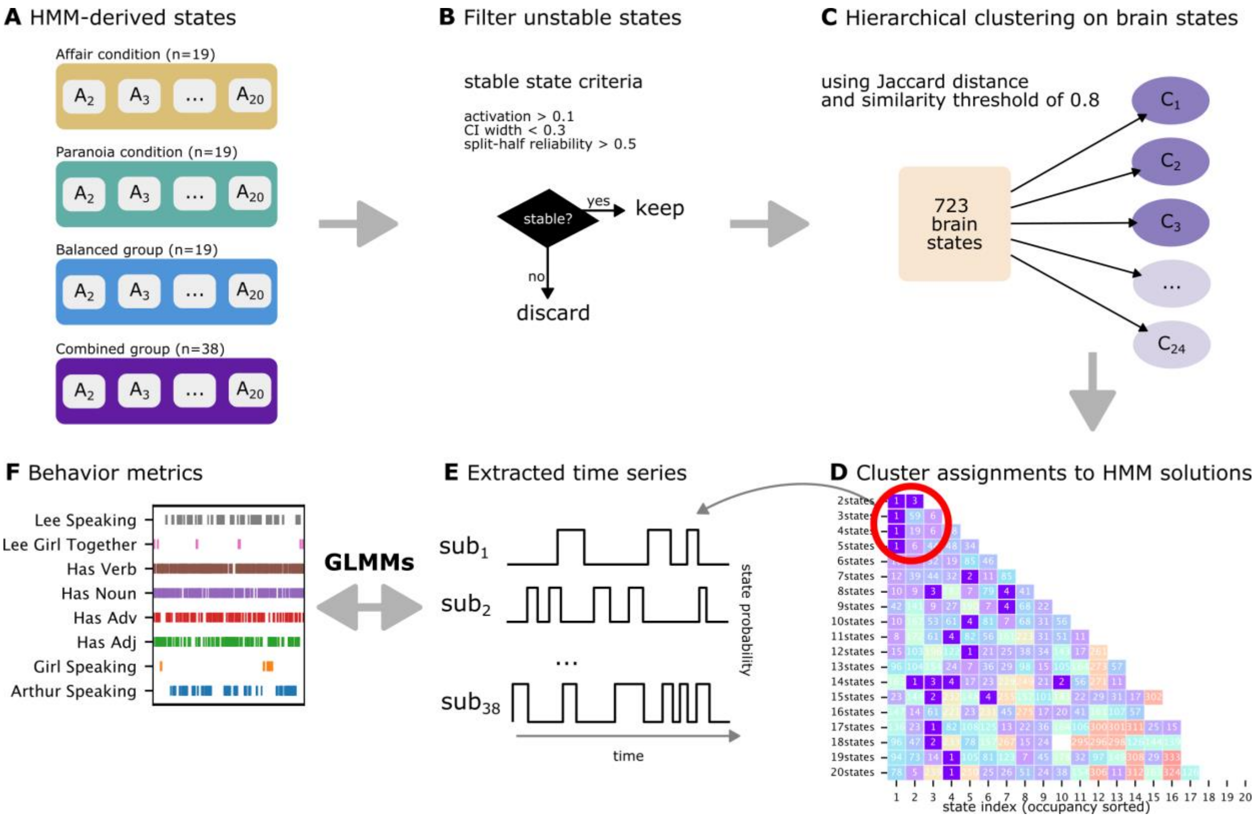


**Supplementary Figure 2. Schematic overview of the analysis pipeline.**

(**A**) Hidden Markov models (HMMs) with 2–20 states were estimated separately for the affair, paranoia, balanced, and combined groups. (**B**) States were retained only if they met stability criteria: activation > 0.1, bootstrap CI width < 0.3, and split-half reliability > 0.5. (**C**) After filtering, 723 reliable states remained. (**D**) These states were grouped using hierarchical clustering with Jaccard distance (similarity threshold = 0.8; thresholds from 0.6–0.9 were tested). This yielded 24 clusters, from which the four largest and most consistent were selected for further analysis. Representative states were traced back to the combined-group HMMs; red circles highlight the first models in which Clusters 1–4 appeared (e.g., Cluster 1: 2-state model; Cluster 2: 7-state model; Cluster 3: 2-state model; Cluster 4: 6-state model). (**E**) For each cluster, we extracted binary time series from the corresponding combined-group HMM, indicating for each participant (n = 38) whether that cluster state was active at each timepoint. (**F**) These binary state time series were then used as predictors in Bayesian generalized linear mixed models (GLMMs) to test associations with linguistic and character annotations from the narrative stimulus.

#### 2. Results

##### 2.1 State pattern clustering

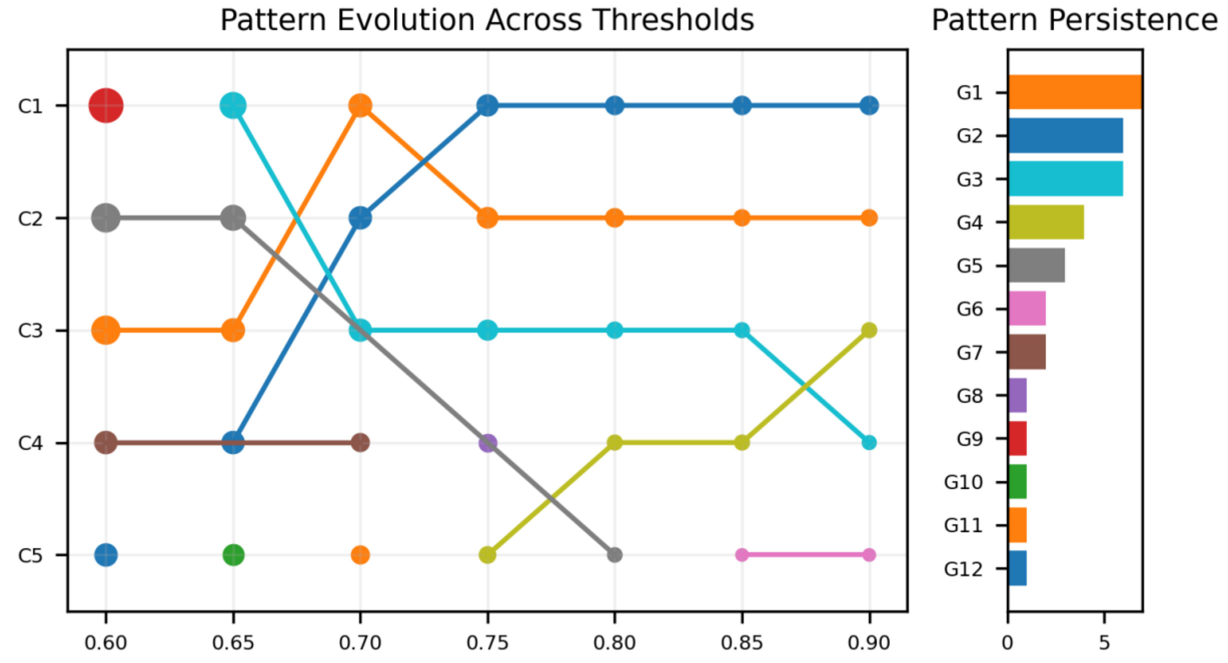


**Supplementary Figure 3. Stability of brain state clusters across similarity thresholds.** (Left) Pattern evolution across thresholds from 0.60 to 0.90. Each line tracks how clusters identified at lower thresholds (C1–C5) persist, merge, or split as the similarity threshold increases, illustrating the stability of cluster identities across a range of cutoffs. (Right) Pattern persistence, quantified as the number of thresholds at which each cluster (G1–G12) was detected, highlights which clusters are most robust to changes in threshold.

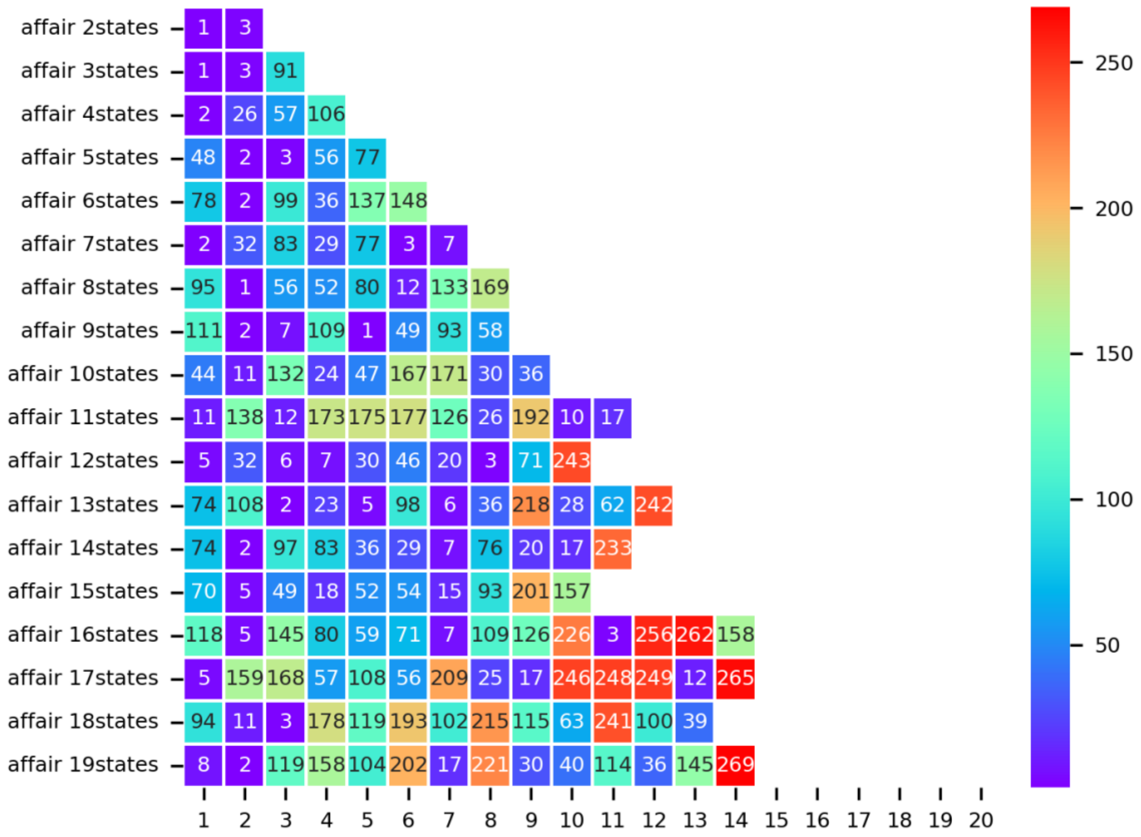


**Supplementary Figure 4. HMM state clustering results for the affair group.**

Each row corresponds to an HMM fit with a different number of states (ranging from 2 to 20), while each column represents a brain state sorted by model-specific occupancy (i.e., the proportion of time spent in each state). Cell colors and overlaid numbers indicate cluster identity. Missing cells indicate states that did not pass the filtering criteria and were therefore excluded from clustering.

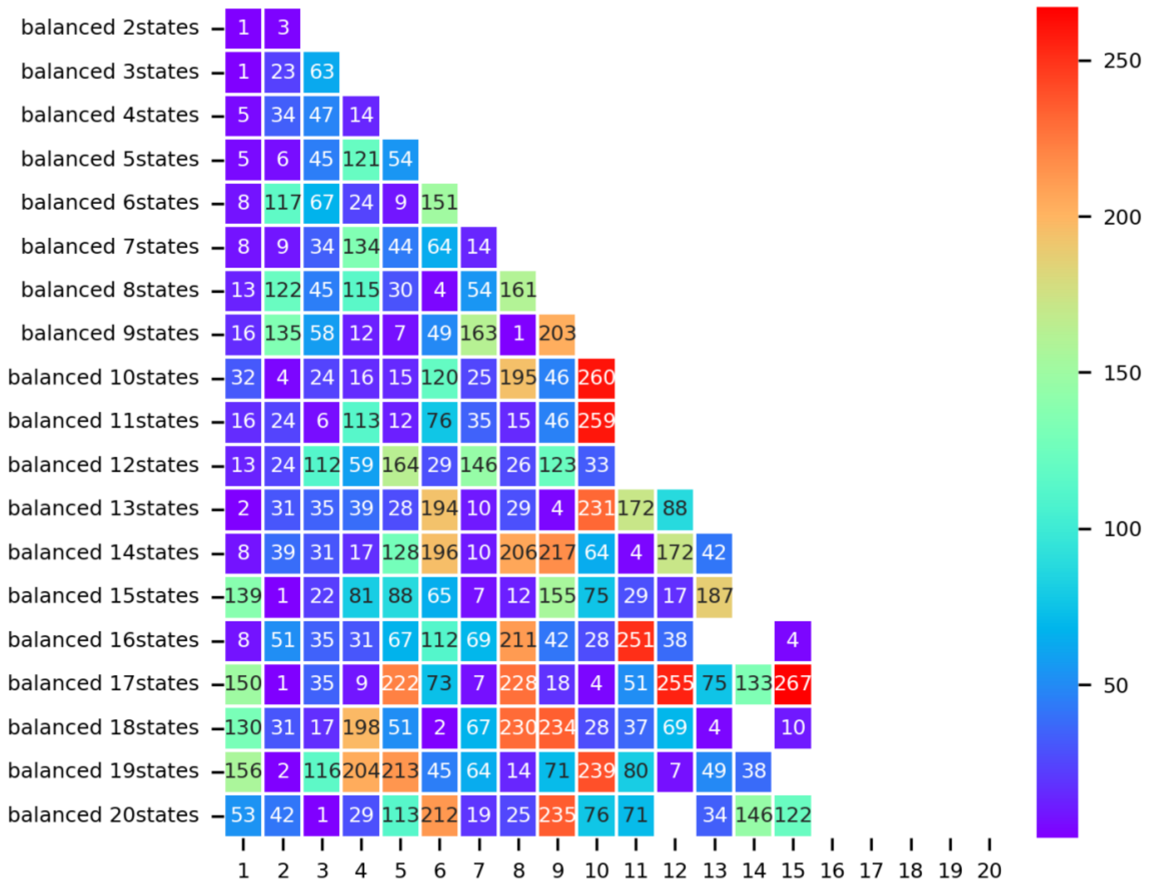


**Supplementary Figure 5. HMM state clustering results for the balanced group.**

Each row corresponds to an HMM fit with a different number of states (ranging from 2 to 20), while each column represents a brain state sorted by model-specific occupancy (i.e., the proportion of time spent in each state). Cell colors and overlaid numbers indicate cluster identity. Missing cells indicate states that did not pass the filtering criteria and were therefore excluded from clustering.

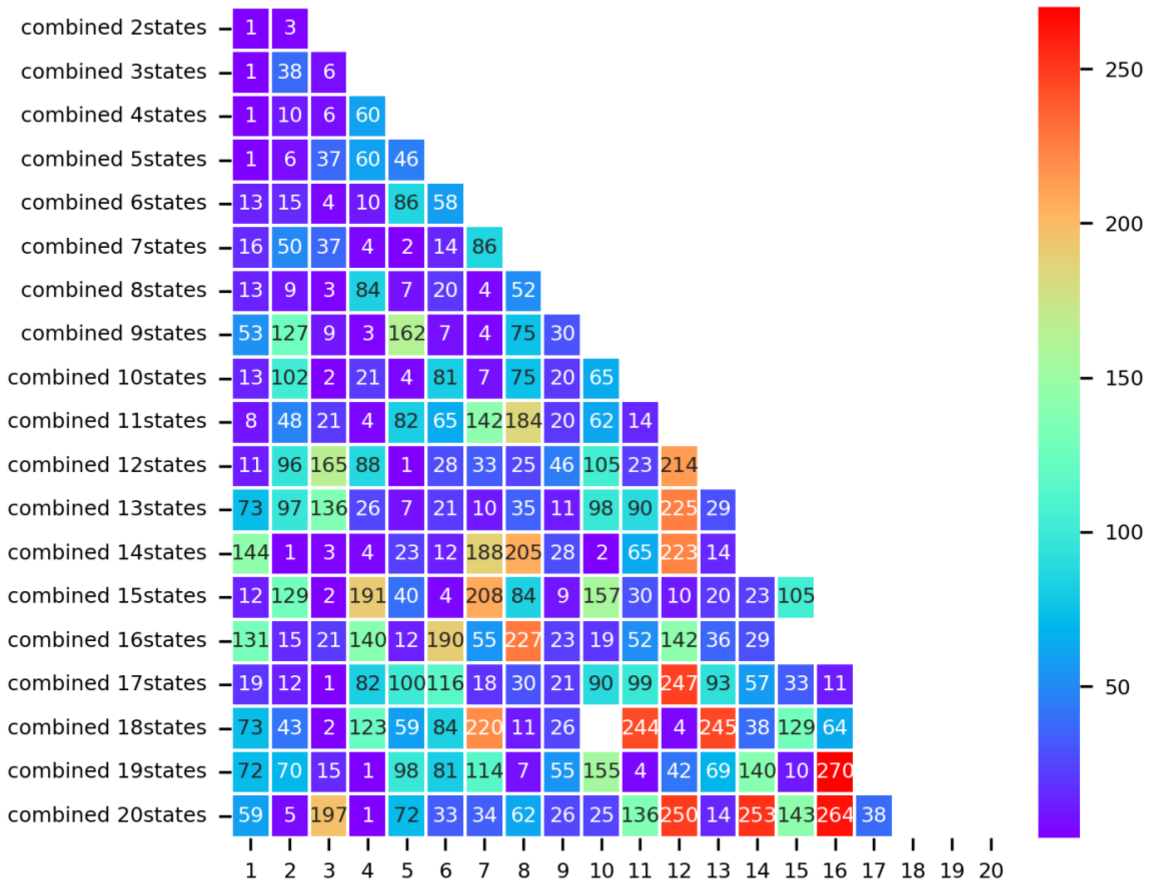

**Supplementary Figure 6. HMM state clustering results for the combined group.**
Each row corresponds to an HMM fit with a different number of states (ranging from 2 to 20), while each column represents a brain state sorted by model-specific occupancy (i.e., the proportion of time spent in each state). Cell colors and overlaid numbers indicate cluster identity. Missing cells indicate states that did not pass the filtering criteria and were therefore excluded from clustering.

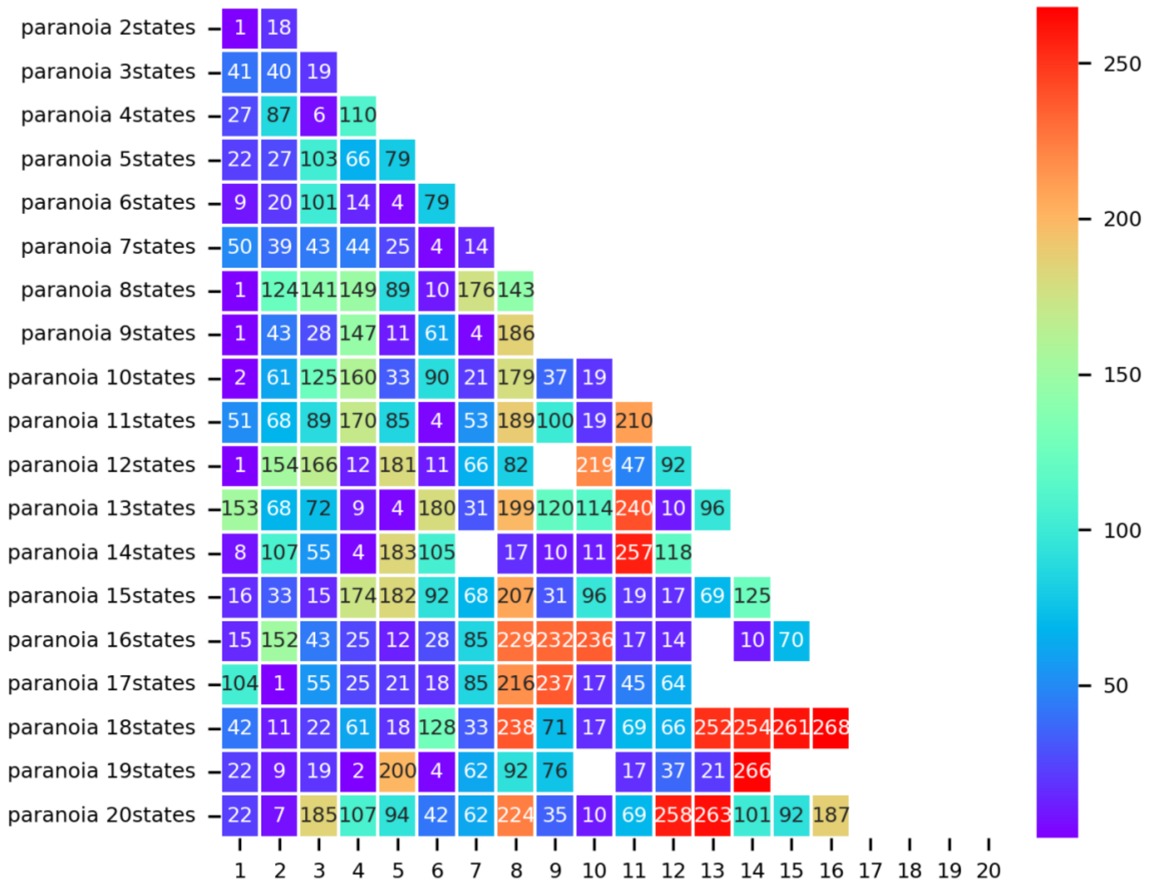


**Supplementary Figure 7. HMM state clustering results for the paranoia group.**

Each row corresponds to an HMM fit with a different number of states (ranging from 2 to 20), while each column represents a brain state sorted by model-specific occupancy (i.e., the proportion of time spent in each state). Cell colors and overlaid numbers indicate cluster identity. Missing cells indicate states that did not pass the filtering criteria and were therefore excluded from clustering.

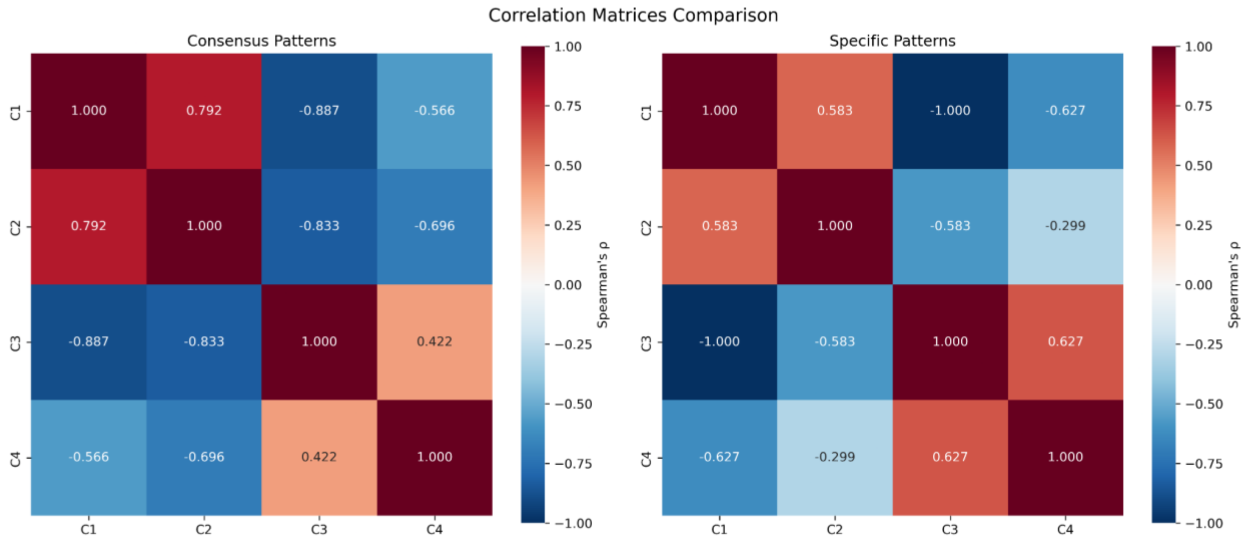


**Supplementary Figure 8. Spearman correlation matrices for brain state clusters.**
Pairwise Spearman rank correlations (ρ) among brain state clusters are shown using two complementary approaches. Left: Consensus patterns, derived by averaging activation vectors across all member states within each cluster. Right: Representative patterns, defined by selecting one exemplar state from the combined-group HMM models for each cluster. Both approaches reveal positive correlations between Clusters 1 and 2 and between Clusters 3 and 4, and negative correlations between the two groups, consistent with qualitative observations. Values indicate correlation coefficients across 17 canonical brain networks.

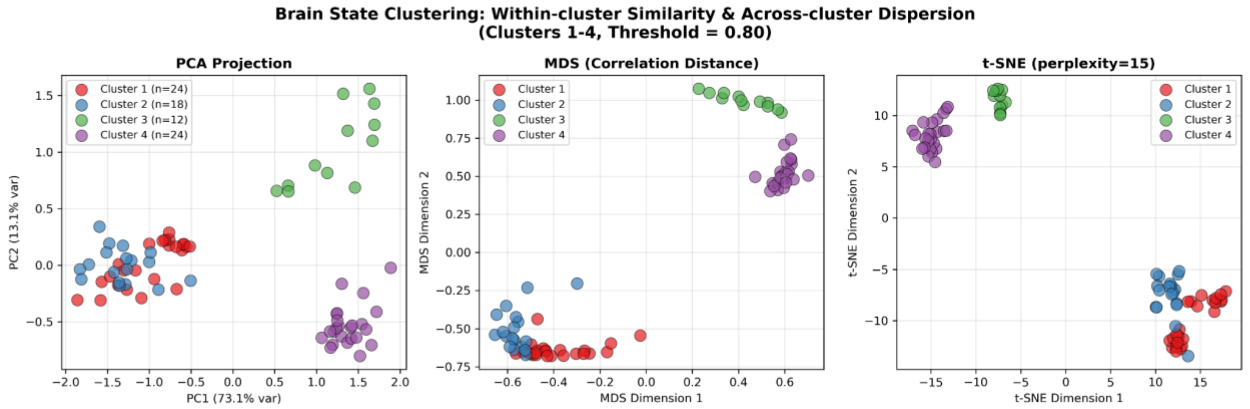


**Supplementary Figure 9. Low-dimensional visualization of clustered brain states.**
Brain state clustering results are shown using three different projection methods: PCA (left), multidimensional scaling (MDS) with correlation distance (middle), and t-SNE with perplexity = 15 (right). Each point corresponds to an individual HMM-derived state pattern, colored by its

assigned cluster (Clusters 1–4). Across all methods, states within the same cluster are tightly grouped, while states from different clusters are well separated, demonstrating high within-cluster similarity and clear across-cluster dispersion.

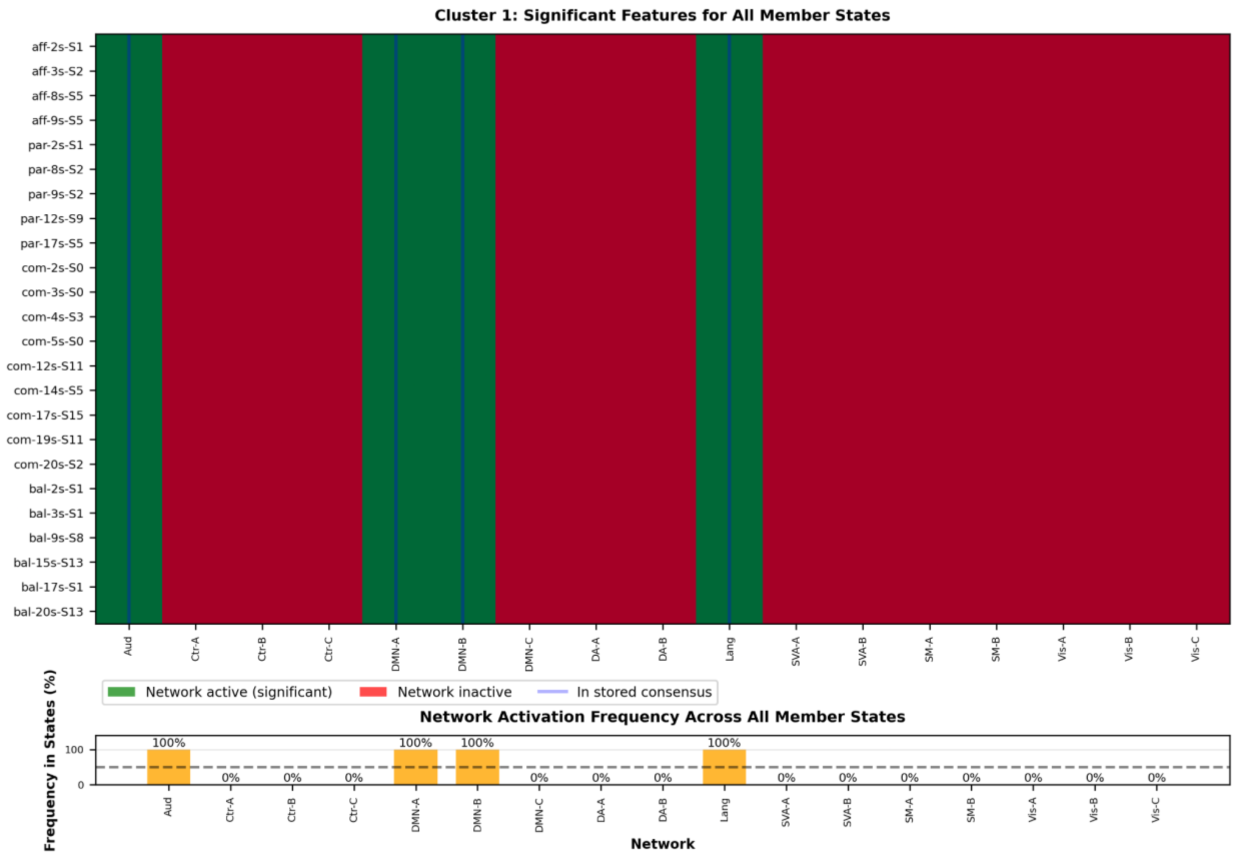


**Supplementary Figure 10. Individual State Composition of Cluster 1**

Cluster 1 represents a canonical narrative comprehension state characterized by coordinated activation of auditory, default mode (DMN-A, DMN-B), and language networks. (A) Heatmap showing network activation patterns for all 24 individual states assigned to Cluster 1, derived from models ranging from 2 to 20 states across all four groups (affair, paranoia, combined, balanced). Green cells indicate networks that met significance criteria (mean activation > 0.1, 95% CI lower bound > 0.05, cross-validation stability ≥ 70%). Blue vertical lines mark networks in the stored consensus pattern. (B) Frequency of each network's activation across all member states. Orange bars show the percentage of states in which each network was significantly active. The black dashed line at 50% indicates the threshold for inclusion in the consensus pattern. All four consensus networks (Aud, DMN-A, DMN-B, Lang) appear in 100% of member states, demonstrating the high stability of this core brain state pattern.

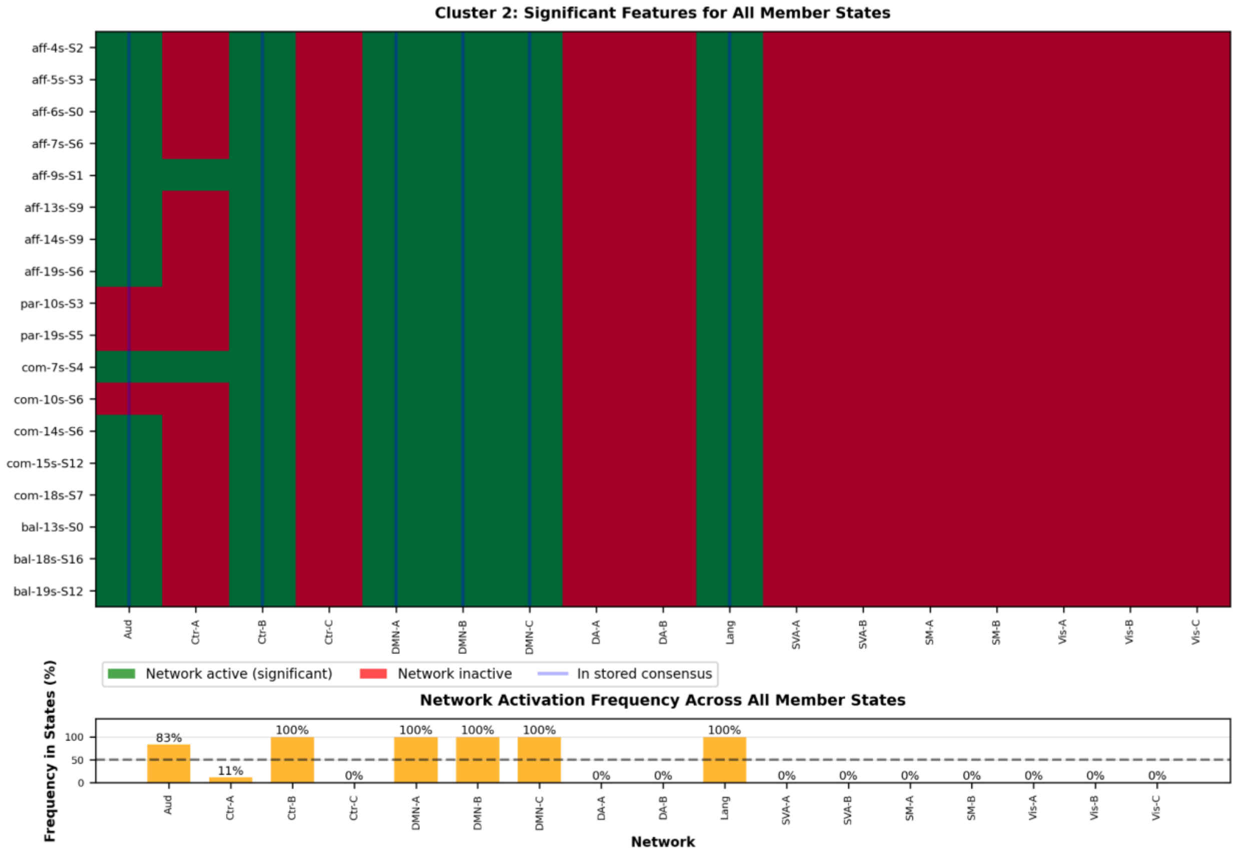


**Supplementary Figure 11. Individual State Composition of Cluster 2**

Cluster 2 represents an elaborative narrative processing state involving auditory, executive control (Ctr-B), default mode (DMN-A, DMN-B, DMN-C), and language networks. (A) Heatmap showing network activation patterns for all 18 individual states assigned to Cluster 2, derived from models ranging from 4 to 19 states across all four groups. Blue vertical lines indicate networks in the stored consensus pattern. (B) Activation frequency across member states. The six consensus networks (Aud, Ctr-B, DMN-A, DMN-B, DMN-C, Lang) appear in 78-100% of states. Additional networks, such as control network A (Ctr-A; present in 11% of states,) appear in specific instances but do not reach the consensus threshold, reflecting context-specific or model-specific variation in how this brain state manifests.

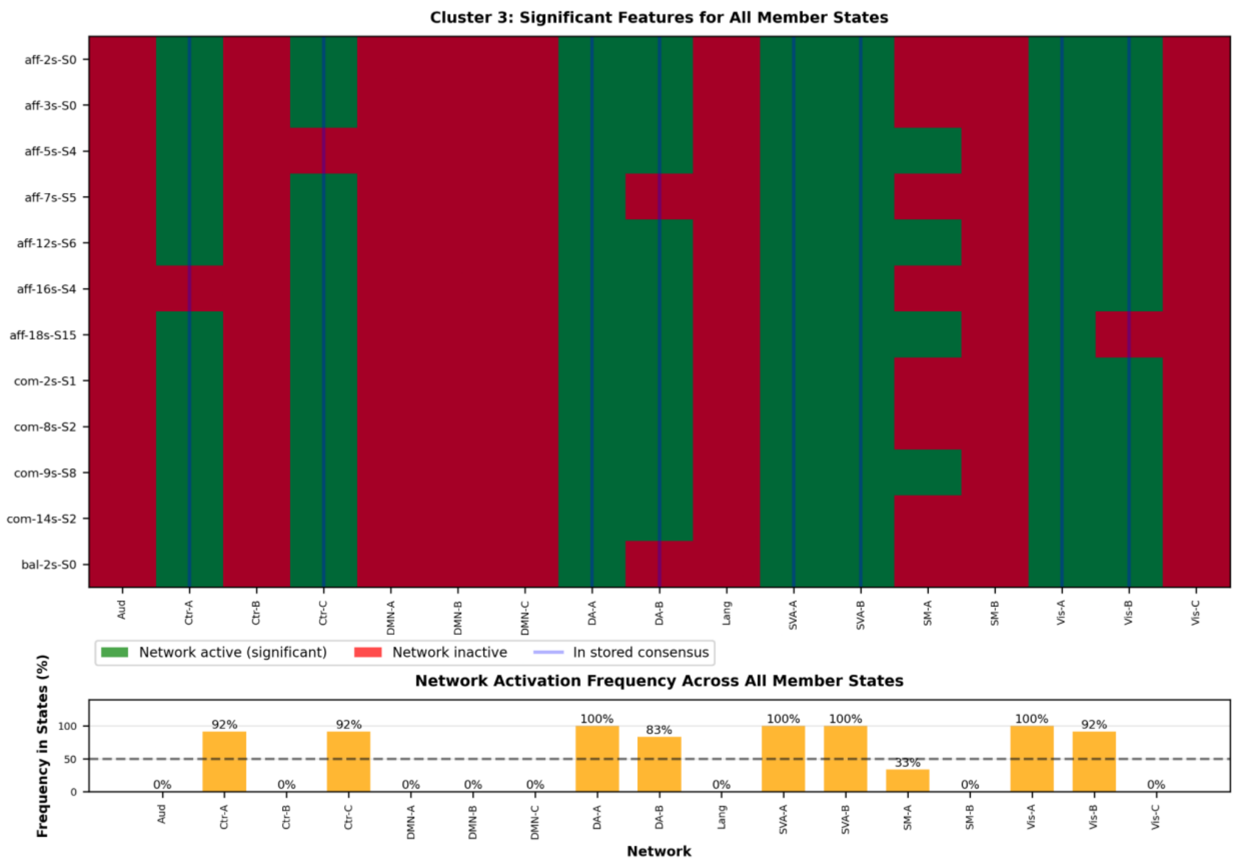


**Supplementary Figure 12. Individual State Composition of Cluster 3**

Cluster 3 represents an externally-oriented perceptual state characterized by activation of sensory and attention networks. (A) Heatmap showing network activation patterns for all 12 individual states assigned to Cluster 3, primarily derived from lower-complexity models (2-18 states) across affair, combined, and balanced groups. Blue vertical lines mark consensus networks. (B) Activation frequency shows that the eight consensus networks (Ctr-A, Ctr-C, DA-A, DA-B, SVA-A, SVA-B, Vis-A, Vis-B) appear in 67-100% of member states. Sensorimotor network A (SM-A) appears in 42% of states, just below the consensus threshold, illustrating how the 50% cutoff identifies the most robust, defining features while accommodating secondary variations.

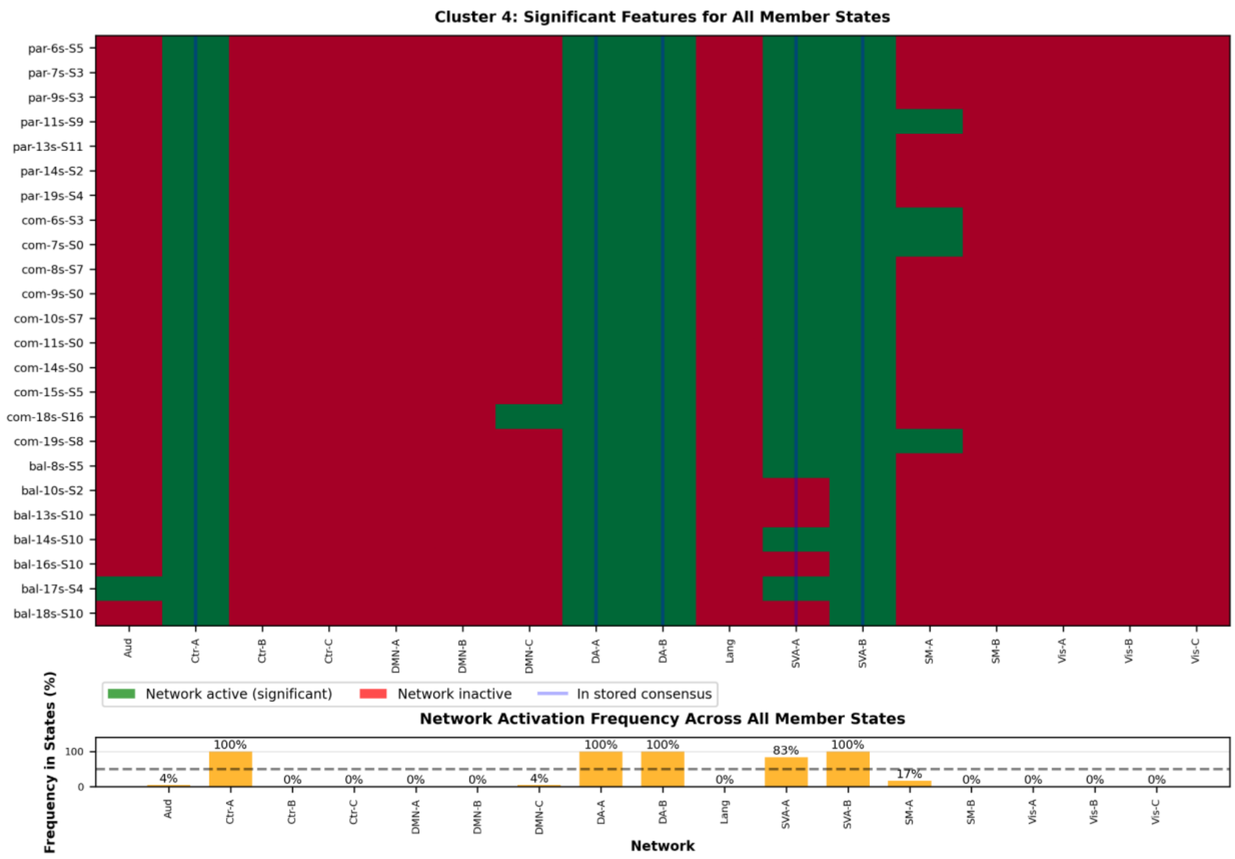


**Supplementary Figure 13. Individual State Composition of Cluster 4**

Cluster 4 represents a focused attention state involving control and dorsal attention networks alongside sensory systems. (A) Heatmap showing network activation patterns for all 24 individual states assigned to Cluster 4, derived from models ranging from 6 to 19 states, with particularly strong representation from the combined and balanced groups. Blue vertical lines indicate consensus networks. (B) Activation frequency demonstrates that the five consensus networks (Ctr-A, DA-A, DA-B, SVA-A, SVA-B) appear in 79-100% of states. Sensorimotor network A (SM-A) appears in 25% of states, representing a subset of instances where this attention state additionally recruits motor preparation systems. This variation illustrates how brain state patterns can exhibit stable core features while flexibly incorporating additional networks depending on specific task demands or narrative contexts.

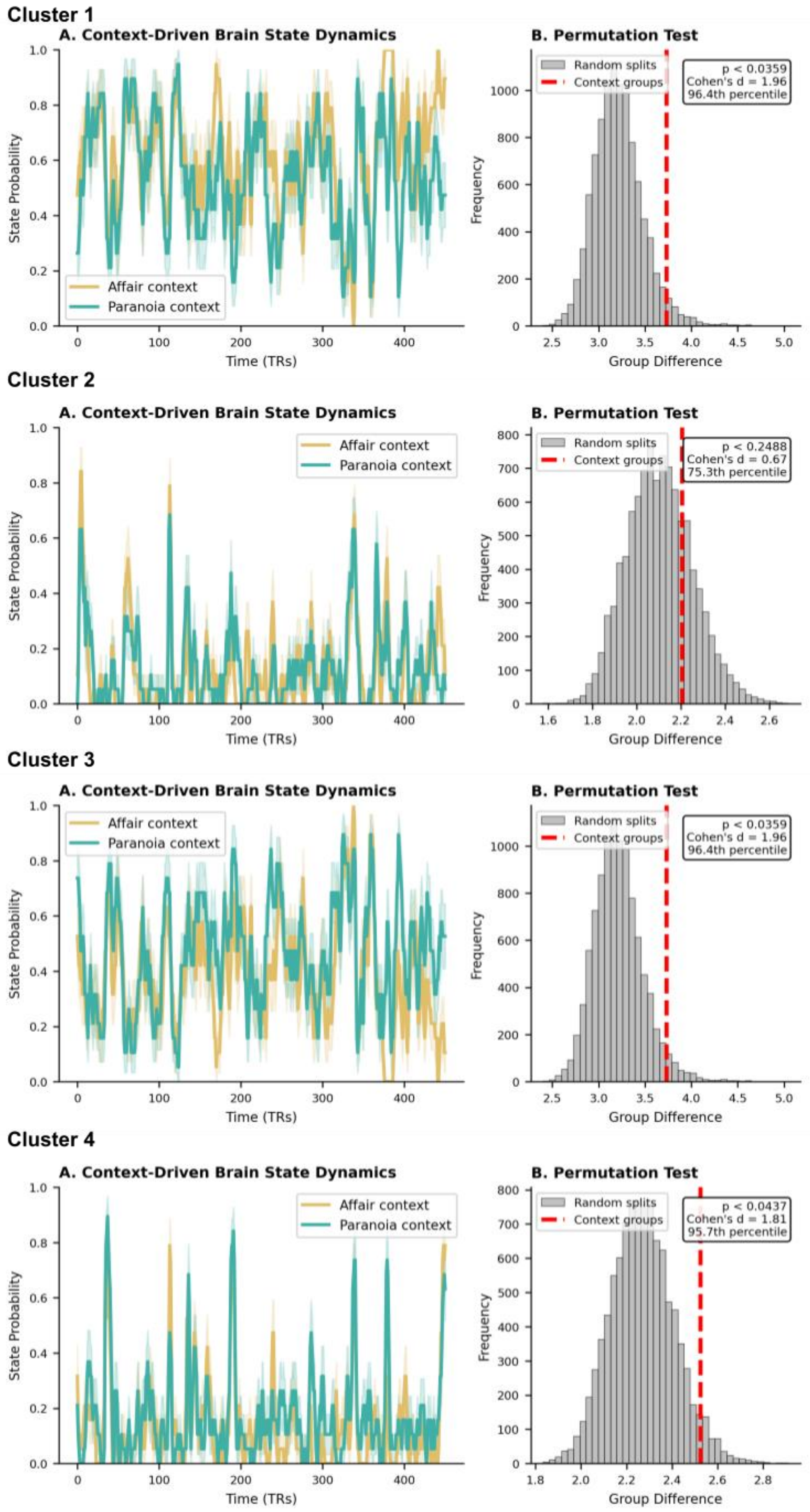


**Supplementary Figure 14. Permutation tests for context-dependent brain state dynamics**

For each of the four clusters, we assessed whether observed group differences in temporal dynamics could arise from arbitrary participant variability. (A) Time series of state probability for affair (gold) and paranoia (teal) context groups, showing moment-to-moment fluctuations in the likelihood of each brain state throughout the narrative. Shaded regions indicate standard error across participants. (B) Permutation test results. Gray histograms show the null distribution of group differences derived from 10,000 random participant splits (irrespective of context assignment). Red dashed lines indicate the observed difference between context groups. For Clusters 1, 3, and 4, the observed context-based differences exceeded the 95th percentile of the null distribution (all $p < .05$, Cohen's $d = 1.81–1.96$), indicating that context grouping produces systematically different state dynamics than would be expected from arbitrary inter-individual variability. Cluster 2 did not show this effect ($p = .249$, Cohen's $d = 0.67$, 75.3rd percentile), suggesting that temporal engagement with this state does not differ reliably between context conditions. Note that Clusters 1 and 3 are derived from the same 2-state combined model and thus have complementary time series (when one is active, the other is not), resulting in identical permutation statistics.

**2.2 Brain state - content GLMM results**

Cluster 1:
https://github.com/yibeichan/prettymouth/blob/main/results/cluster1_combined_2states_deviation_th080.md

Cluster 2:
https://github.com/yibeichan/prettymouth/blob/main/results/cluster2_combined_7states_deviation_th080.md

Cluster 3:
https://github.com/yibeichan/prettymouth/blob/main/results/cluster3_combined_2states_deviation_th080.md

Cluster 4:
https://github.com/yibeichan/prettymouth/blob/main/results/cluster4_combined_6states_deviation_th080.md

**2.3 Behavior - content GLMM results**

Bin at TR (1.5s):

- Figure: https://github.com/yibeichan/prettymouth/blob/main/notebooks/figure5.ipynb
- Results: https://github.com/yibeichan/prettymouth/blob/main/results/behavioral_content_analysis.md

Bin at 1s:

- Figure: https://github.com/yibeichan/prettymouth/blob/main/notebooks/RR_figure5_supplement.ipynb
- Results: https://github.com/yibeichan/prettymouth/blob/main/results/behavioral_content_analysis_supplement.md
